# Genome Profiling for Aflatoxin B1 Resistance in *Saccharomyces cerevisiae* Reveals a Role for the CSM2/SHU Complex in Tolerance of Aflatoxin B_1_-associated DNA Damage

**DOI:** 10.1101/629436

**Authors:** Nick St. John, Julian Freedland, Henri Baldino, Frank Doyle, Cinzia Cera, Thomas Begley, Michael Fasullo

**Author notes:** RNA Institute, State University of New York at Albany, 1400 Washington Ave, Albany, New York 12222. Public Depository of Data: https://www.ncbi.nlm.nih.gov/geo/query/acc.cgi?acc=GSE129699. Corresponding author: Michael Fasullo, College of Nanoscale Science and Engineering, State University of New York Polytechnic Institute, 257 Fuller Drive, Albany, New York 12203.

## Abstract

Exposure to the mycotoxin aflatoxin B1 (AFB_1_) strongly correlates with hepatocellular carcinoma. P450 enzymes convert AFB_1_ into a highly reactive epoxide that forms unstable 8,9-dihydro-8-(*N*7-guanyl)-9-hydroxyaflatoxin B1 (AFB_1_-*N*^7^-Gua) DNA adducts, which convert to stable mutagenic AFB_1_ formamidopyrimidine (FAPY) DNA adducts. In CYP1A2-expressing budding yeast, AFB_1_ is a weak mutagen but a potent recombinagen. However, few genes have been identified that confer AFB_1_ resistance. Here, we profiled the yeast genome for AFB_1_ resistance. We introduced the human CYP1A2 into ∼90% of the diploid deletion library, and pooled samples from CYP1A2-expressing libraries and the original library were exposed to 50 μM AFB_1_ for 20 hs. By using next generation sequencing to count molecular barcodes, we identified 85 AFB_1_ resistant genes from the CYP1A2-expressing libraries. While functionally diverse genes, including those that function in proteolysis, actin reorganization, and tRNA modification, were identified, those that function in post-replication DNA repair and encode proteins that bind to DNA damage were over-represented, compared to the yeast genome, at large. DNA metabolism genes included those functioning in DNA damage tolerance, checkpoint recovery and replication fork maintenance, emphasizing the potency of the mycotoxin to trigger replication stress. Among genes involved in error-free DNA damage tolerance, we observed that *CSM2*, a member of the *CSM2(SHU)* complex, functioned in AFB_1_-associated sister chromatid recombination while suppressing AFB_1_-associated mutations. These studies thus broaden the number of AFB_1_ resistant genes and have elucidated a mechanism of error-free bypass of AFB_1_-associated DNA adducts.

## INTRODUCTION

The mycotoxin aflatoxin B1 (AFB_1_) is a potent hepatocarcinogen. The signature p53 mutation, p53-Ser249, is often present in liver cancer cells from hepatocellular carcinoma (HCC) patients from AFB_1_-exposed areas, suggesting that AFB_1_ is a potent carcinogen because it is a genotoxin (Hsu et al., 1991; Shen and Ong, 1996). A mutagenic signature associated with AFB_1_ exposure has been identified in HCC (Chawanthayatham et al., 2017; Huang et al., 2017). However, AFB_1_ is not genotoxic *per se* but is metabolically activated by P450 enzymes, such as CYP1A2 and CYP3A4 (Crespi et al., 1991; Eaton and Gallagher, 1994; Gallagher et al., 1996), to form a highly reactive AFB_1_-8-9-epoxide (Baertschi et al., 1988). The epoxide reacts with protein, RNA, and DNA, yielding the unstable 8,9-dihydro-8-(*N*7-guanyl)-9-hydroxyaflatoxin B1 (AFB1-*N*^7^-Gua) adducts that convert into stable AFB_1_-formamidopyrimidine (FAPY) adducts (Essigman et al., 1977; Lin et al., 1977; Croy et al., 1981). The anomers of the AFB_1_-FAPY-DNA adduct block DNA replication or cause mutations in *Escherichia coli* (Smela et al., 2002; Brown et al., 2006) and *in vitro* (Lin et al, 2014). Metabolically active AFB_1_ can also indirectly damage DNA through oxidative stress (Shen et al., 1995; Bedard and Masey, 2006; Bernabucci et al., 2011; Singh et al., 2015). Identifying genes that repair AFB_1_-associated DNA damage could help identify which individuals are at elevated risk for HCC. However, epidemiological data has been inconsistent, and only a few candidate DNA repair genes have been proposed, such as XRCC1(Pan et al., 2011, Xu et al., 2015), XRCC3 (Long et al., 2008; De Mattia et al., 2017) and XRCC4 (Long et al., 2013).

AFB_1_ resistance genes have been identified from model organisms, revealing mechanisms by which AFB_1_-associated DNA adducts can be both tolerated and excised. Both prokaryotic and eukaryotic nucleotide excision repair (NER) pathways function to remove AFB_1_-associated DNA adducts (Leadon, et al, 1981; Alekseyev et al., 2004; Bedard and Massey, 2006). Recently, the base excision repair gene (BER) NEIL1 has been implicated in direct repair AFB_1_-associated DNA adducts (Vartanian et al., 2017). To tolerate persistent AFB_1_-associated DNA lesions, translesion polymerases in yeast, such as those encoded by *REV1* and *REV7*, and in mouse, such as Polζ, confer resistance and may promote genome stability (Lin et al., 2014).

While yeast do not contain endogenous P450 genes that can metabolically activate AFB_1_ into an active genotoxin (Sengstag et al., 1996; Van Leeuwen et al., 2013, Fasullo et al., 2014), human CYP1A2 can be expressed in yeast cells (Sengstag et al., 1996; Fasullo et al., 2014). Interestingly, metabolically activated AFB_1_ is a potent recombinagen but weak mutagen (Sengstag et al., 1996). CYP1A2-activated AFB_1_ reacts to form both the unstable AFB_1_-N^7^-Gua adducts and the stable AFB_1_-FAPY DNA adducts (Fasullo et al., 2008). The AFB_1_-associated DNA damage, in turn, triggers a robust DNA damage response that includes checkpoint activation (Fasullo et al., 2008), cell cycle delay (Fasullo et al., 2010), and the transcriptional induction of stress-induced genes (Keller-Seitz et al., 2004; Guo et al., 2006). Profiles of the transcriptional response to AFB_1_ exposure reveals induction of genes in growth and checkpoint signaling pathways, DNA and RNA metabolism, and protein trafficking (Keller-Seitz et al, 2004; Guo et al., 2006). While genes involved in recombinational repair and DNA damage tolerance confer AFB_1_ resistance (Keller-Seitz et al., 2004; Fasullo et al, 2010; Guo et al, 2005), it is unclear the functional significance of many genes in the stress induced pathways in conferring resistance since transcriptional induction is not synonymous with conferring resistance (Birrell et al., 2004).

In this study, we profiled the yeast genome for AFB_1_ resistance. We asked which genes confer AFB_1_ resistance in the presence or absence of human CYP1A2 expression by screening the non-essential diploid collection by high throughput sequencing of molecular barcodes (Pierce et al., 2006; St Onge et al. 2007; Smith et al., 2010). While we expected to identify NER, recombinational repair, and DNA damage tolerance genes, which had previously been identified (Keller-Seitz et al., 2004; Fasullo et al, 2010; Guo et al, 2005), our high throughput screen identified novel genes involved in AFB_1_ resistance. These included genes involved in Rad51 assembly, cell cycle progression, RNA metabolism, and oxidative stress. Our results thus underscore the importance of recombination in both mutation avoidance and in conferring AFB_1_ resistance.

## Materials and Methods

### Strains and plasmids

Yeast strains were derived from BY4741, BY4743 (Brachman et al., 1998) or YB204 (Dong and Fasullo, 2003); all of which are of the S288C background (supplemental Table 1). The BY4743 genotype is *MAT*a/α *his3*Δ*1/his3*Δ*1 leu2*Δ*0/leu2*Δ*0 LYS2/lys2*Δ*0 met15*Δ*0/MET15 ura3*Δ*0/ura3*Δ*0.* The diploid and haploid homozygous deletion libraries were purchased from Open Biosystems, and are now available from Dharmacon (http://dharmacon.gelifesciences.com/cdnas-and-orfs/non-mammalian-cdnas-and-orfs/yeast/yeast-knockout-collection/). The pooled diploid homozygous deletion library (n = 4607) was a gift of Chris Vulpe (University of Florida).

To construct the *csm2 rad4* and *csm2 rad51* double mutants, we first obtained from haploid *csm2* strain (YA288, supplemental Table 1) from the haploid BY4741-derived deletion library. We introduced the *his3* recombination substrates (Fasullo and Davis, 1987) to measure unequal sister chromatid exchange (SCE) in the *csm2* mutant by isolating the meiotic segregant YB558 from a diploid cross of YB204 with YA288. This *csm2* strain (YB558) was subsequently crossed with *MAT*α *rad4::NatMX* (YA289) and the *csm2::KanMX rad4::NatMX* meiotic segregant (YB600) was obtained. The *rad51 csm2* double mutant was made by one step gene disruption (Rothstein, 1983) using the *Bam*H1 fragment *rad51*Δ (Shinohara et al., 1992) to select for Ura^+^ transformants in YB558.

Using LiAc-mediated gene transformation we introduced human CYP1A2 into BY4741, *csm2*, *rad4*, *rad51*, *csm2 rad51*, and *csm2 rad4* strains. The CYP1A2-expression plasmid, pCS316, was obtained by CsCl centrifugation (Ausubel, 1994) and the restriction map was verified based on the nucleotide sequence of the entire plasmid. An alternative CYP1A2-expression plasmid, pCYP1A2-NAT2 was constructed by removing the hOR sequence from pCS316 and replacing it with a *Not*1 fragment containing the human NAT2.

### Media and chemicals

Standard media were used for the culture of yeast and bacterial strains (Burke et al., 2000). LB-AMP (Luria broth containing 100 μg/ml ampicillin) was used for the culture of the bacterial strain DH1 strain containing the vector pCS316. Media used for the culture of yeast cells included YPD (yeast extract, peptone, dextrose), SC (synthetic complete, dextrose), SC-HIS (SC lacking histidine), SC-URA (SC lacking uracil), and SC-ARG (SC-lacking arginine). Media to select for canavanine resistance contained SC-ARG (synthetic complete lacking arginine) and 60 μg/mL canavanine (CAN) sulfate, and media to select for 5-fluoroorotic acid (FOA) resistance contained SC-URA supplemented with 4x uracil and FOA (750 μg/ml), as described by Burke et al. (2000). FOA plates contained 2.2% agar; all other plates contained 2% agar. AFB_1_ was purchased from Sigma Co., and a 10 mM solution was made in dimethyl sulfoxide (DMSO).

#### Measuring DNA Damage-Associated Recombination and Mutation Events

To measure AFB_1_-associated genotoxic events, log phase yeast cells (*A*_600_=0.5-1) were exposed to indicated doses of AFB_1_, previously dissolved in DMSO. Cells were maintained in synthetic medium (SC-URA) during the carcinogen exposure. After the exposure, cells were washed twice in H_2_O, and then plated on SC-HIS or SC-ARG CAN to measure unequal SCE or mutation frequency, respectively. An appropriate dilution was inoculated on YPD to measure viability (Fasullo et al., 2008).

### Construction of CYP1A2-expressiong library

To introduce CYP1A2 (pCS316, Sengstag et al., 1996, and pCYP1A2_NAT2) into the yeast diploid deletion collection, we used a modified protocol for high throughput yeast transformation in 96-well plates (Gietz and Schiestl 2006). In brief, FOA^R^ isolates were isolated from each individual strain in the diploid collection and inoculated in 96-well plates, each containing 100 µl of YPD medium. After incubation over-night at 30°C, plates were centrifuged, washed in sterile H_2_O, and resuspended in one-step buffer (0.2 N LiAc, 100 mM DTT, 50% PEG, MW 3300, 500 µg/ml denatured salmon sperm DNA). After addition of 1 µg pCS316 and incubation for 30 minutes at 30°C, 10 µl were directly inoculated on duplicate SC-URA plates. Two Ura^+^ transformants were chosen corresponding to each well and frozen in SC-URA 0.75% DMSO. We introduced the CYP1A2-containing plasmids into approximately 90% of the deletion collection.

### Functional profiling of the yeast genome

The CYP1A2-expressing libraries were pooled and frozen in SC-URA medium containing 0.75% DMSO (n = 4150). The pooled cells (100 µl) were added to 2 ml of SC-URA and allowed to recover for two hours. Cell were then diluted to A_600_ = 0.85 in 2 ml of SC-URA and exposed to either 50 µM AFB_1_ in 0.5% DMSO, and 0.5% DMSO alone. Cells were then incubated with agitation at 30°C for 20 hs. Similarly, the pooled BY4743 library (n = 4607) was directly diluted to A_600_ = 0.85 in YPD and also exposed to 50 µM AFB_1_ and DMSO for 20 hs. Independent triplicate experiments were performed for each library and each chemical treatment.

After AFB_1_ exposure, cells were washed twice in sterile H_2_O and frozen at −80°C. Cells were resuspended in 10 mM Tris-HCl, 1 mM EDTA, 100 mM NaCl, 2% Triton X-100, 1% SDS, pH 8 and DNA was isolated by “smash and grab (Hoffman and Winston. 1987).” Barcode sequences, which are unique for each strain in the deletion collection (Giaever et al., 2002; Giaever et al, 2004), were amplified by PCR using a protocol described by Smith et al. (2010). The primers used for amplification are listed in the Supplemental Table 2. 125 bp PCR products were then isolated from 10% polyacrylamide gels by diffusion in 0.5M NH_4_Ac 1 mM EDTA for 24 hs (30°C) followed by ethanol precipitation. The DNA was quantified after being resuspended in Tris EDTA pH 7.5 and the integrity of the DNA was verified by electrophoresis on 10% polyacrylamide. Equal amounts of DNA were pooled from treated and untreated samples. The uptags were then sequenced using the Illumina Platform at the University Buffalo Genomics and Bioinformatics Core (Buffalo, New York). Sequence information was then uploaded to an accessible computer server for further analysis. The software to demultiplex the sequence information, match the uptag sequences with the publish ORFs, and calculate the statistical significance of the differences in log2N ratios was provided by F. Doyle. Tag counts were analyzed with the TCC Bioconductor package (Sun et al., 2013) using TMM normalization (Robinson and Oshlock, 2010) and the edgeR test method (Robinson et al., 2009). Statistical significance was determined using Linux software. Data files have been deposited in the Gene Expression Omnibus database, GSE129699.

### Over-enrichment analysis

Gene Ontology (GO) categories were identified by a hypergeometric distribution with freely available software from University of Toronto (http://funspec.med.utoronto.ca/), YeastMine database (http://yeastmine.yeastgenome.org/), and Panther (http://pantherdb.org/tools/) with a P value cutoff of < 0.05 (Cherry et al., 2012, Mi et al., 2016). The AFB_1_ sensitivity of selected of mutants corresponding to gene ontology groups were verified by growth curves.

### Western blot analysis

Expression of CYP1A2 was determined by Western blots and MROD assays. Cells were inoculated in SC-URA medium. Cells in log growth phase (A_600_ = 0.5–1) were concentrated and protein extracts were prepared as previously described by Foiani et al. (1994). Proteins were separated on 10% acrylamide/0.266% bis-acrylamide gels and transferred to nitrocellulose membranes. Human CYP1A2 was detected by Western blots using goat anti-CYP1A2 (Abcam), and a secondary bovine anti-goat antibody. For a loading control on Western blots, β-actin was detected using a mouse anti-β-actin antibody (Abcam 8224) and a secondary goat anti-mouse antibody. Signal was detected by chemiluminescence, as in previous publications (Fasullo et al., 2014).

### Measuring CYP1A2 enzymatic activity

We measured CYP1A2 enzymatic activity using a modified protocol described by Pompon *et al*. (1996). In brief, cells obtained from 100 ml of selective media were pelleted and resuspended in 5 ml Tris EDTA KCl (pH 7.5, TEK) buffer. After five minute incubation at room temperature, cells were pelleted, resuspended in 1 ml 0.6 M Sorbitol Tris pH 7.5, and glass beads were added. Cells were lysed by agitation. The debris was pelleted at 10,000 x g at 4°C, and the supernatant was diluted in 0.6 M Sorbitol Tris pH 7.5 and made 0.15 M in NaCl and 1% in polyethylene glycol (MW 3350) in a total volume of 5 ml. After incubation on ice for 1 hr. and centrifugation at 10,000 rpm for 20 minutes, the precipitate was resuspended in Tris 10% glycerol pH 7.5, and stored at ^-^80°C.

CYP1A2 enzymatic activity was measured in cell lysates by quantifying 7-methoxyresorufin O-demethylase (MROD) activities (Fasullo et al., 2014), using a protocol similar to that quantifying ethoxyresorufin O-deethylase (EROD) activity (Eugster et al., 1992; Sengstag et al., 1994). The buffer contained 10 mM Tris pH 7.4, 5µM methoxyresorufin (Sigma) and 500 µM NADPH. The production of resorufin was measured in real-time by fluorescence in a Tecan plate reader, calibrated at 535 nm for excitation and 580 nm for absorption, and standardized using serial dilutions of resorufin. The reaction was started by the addition of NADPH and resorufin was measured at one minute intervals during the one hour incubation at 37°C; rat liver microsomes (S9) were used as a positive control while the reaction without NADPH served as the negative control. Enzyme activities were measured in duplicate for at least two independent lysates from each strain and expressed in pmol/min/mg protein.

### Growth assays in 96 well plate to measure AFB_1_ sensitivity

In brief, individual saturated cultures were prepared for each yeast strain. Cell density was adjusted to ∼0.8 x 10^7^ cells/ml for all cultures. We maintained the cells in selective medium (SC-URA). In each microtiter well, 95 µl of media and 5 µl of cells (8 x 10^4^ cells) were aliquoted in duplicate for blank, control and experimental samples. For experimental samples, we added AFB_1_, dissolved in DMSO, for a final concentration of 50 µM and 100 µM. The microtiter dish was placed in a plate reader that is capable of both agitating and incubating the plate at 30°C, as previously described (Fasullo et al., 2010; Fasullo et al., 2014). We measured the A_600_ at 10 minute intervals, for a total period for 24 hs, 145 readings. Data at 1hr intervals was then plotted. To avoid evaporation during the incubation, the microtiter dishes were sealed with clear optical tape (Fasullo et al, 2010). To calculate area under the curve (AUC), we used a free graphing application (https://www.padowan.dk/download/), and measured the time interval between 0-20 hs, as performed in previous publications (O’Connor et al., 2012). Statistical significance was determined by the Student’s *t*-test, assuming constant variance between samples.

To determine epistasis of AFB_1_-resistant genes, we calculated the deviation ε according to ε = W_xy_-W_x_x W_y_, where W_x_ and W_y_ are the fitness coefficients determined for each single mutant exposed to AFB_1_ and W_xy_is the product. Fitness was calculated by determining the generation time of both the single and double mutants over three doubling times. Zero and negative values are indicative of genes that do not interact or participate in the same pathway to confer fitness (St. Onge et al., 2007).

### Data Availability

All yeast strains and plasmids are available upon request and are detailed in STable 1. Three supplementary tables and two supplementary figures have been deposited in figshare. Next generation sequencing data (NGS) of barcodes are available at GEO with accession number GSE129699.

## Results

We used three BY4743-derived libraries to profile the yeast genome for AFB_1_ resistance. The first was a pooled library of 4607 yeast strains, each strain containing a single deletion in a non-essential gene (Jo et al, 2009). The second was a pooled library of approximately 4900 strains each containing individual deletions in non-essential genes and was made by introducing pCS316 into each strain by yeast transformation (Figure 1). The third was a pooled library of approximately 5000 strains expressing both CYP1A2 and NAT2. By calculating area under the growth curves, we determined that the D_10_ for wild type BY4743 expressing CYP1A2 was 50 μM AFB_1_ while the D_16_ for BY4743 expressing CYP1A2 was 100 μM (Figure 1). The D_10_ for wild type BY4743 expressing CYP1A2 with the hOR was the same as in BY4743 cells expressing CYP1A2 and NAT2. To confirm metabolic activation of AFB_1_ into a potent genotoxin, we showed that growth of the *rad52* diploid mutant was significantly impaired (Figure 1). The non-linear relationship between dose and lethality is consistent with previous results that the number of DNA adducts do not linearly increase with increasing AFB_1_ concentrations (Fasullo et al., 2008). Cells that did not express CYP1A2 showed slight growth delay after cells were exposed to 100 µM AFB_1_ (Figure 1).

**Figure 1:**
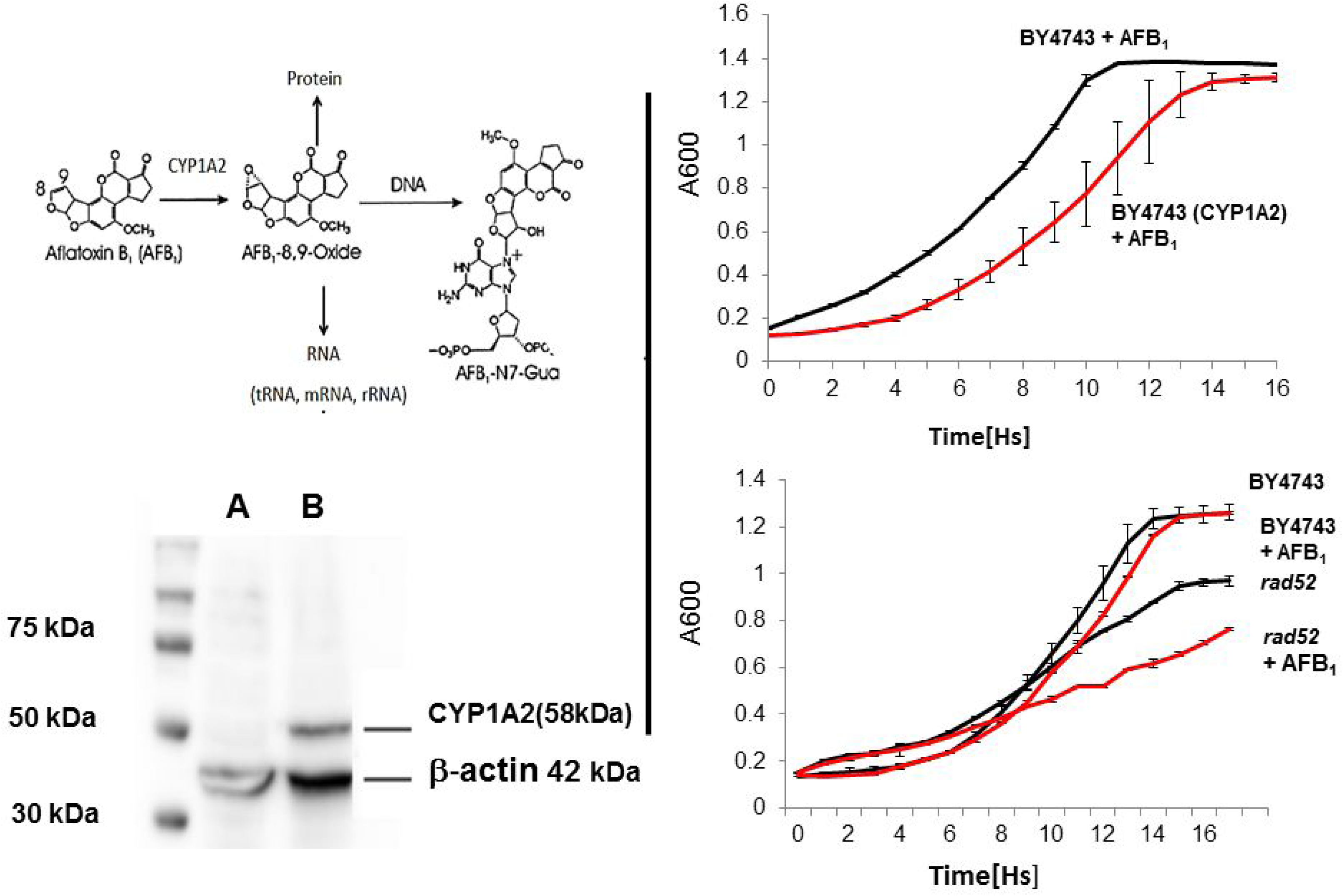
Expression of CYP1A2 in the yeast diploid strain (BY4743). Top right indicates CYP1A2-mediated activation of AFB_1_ to form a highly reactive epoxide that forms DNA, RNA, and protein adduct. The lower left panel is a Western blot indicating 75, 50 and 37 kDa molecular weight markers. Lanes A and B are lysates from BY4743 and BY4743 cells expressing CYP1A2, respectively. The CYP1A2 (58 kDa) protein and the β-actin (42 kDa) protein are indicated. Right upper panel is a growth curve of the diploid wild type (BY4743) and BY4743 + pCS316 after exposure to 100 µM AFB. Right lower panel is a growth curve of BY4743 + pCS316 and *rad52* + pCS316 after exposure to 1% DMSO and 100 µM AFB_1_. Growth (A_600_) is plotted against time (Hs). Standard deviations are indicated at 1 h time points.

### Confirmation of CYP1A2 activity

To confirm that CYP1A2 was both present and functional in the library, we performed Western blots and MROD assays, as in previous studies (Fasullo et al., 2014; Figure 1). Two independent assays were performed for five different ORFs (*RAD2, RAD18, RAD55, OGG1, MRC1*); the average MROD result was 5-10 units pmol/sec/mg protein. These results are similar to what was observed for the wild type BY4743 (Fasullo et al., 2014) and for various haploid mutants (Guo et al., 2005). These studies indicate that CYP1A2 is active in diploid strains and can be detected by Western blots in BY4743-derived strains containing pCS316, in agreement with previous studies (Guo et al., 2005).

### Identification of Genes by Barcode Analysis

We initially grouped AFB_1_ resistance genes into: 1) those that confer resistance to AFB_1_ without CYP activation, and 2) those that confer resistance to P450-activated AFB_1_. After exposing cells to 50 µM AFB_1_, we identified barcodes for approximately 51% and 89% of the genes from pooled deletion library with and without CYP1A2, respectively (for complete listing, see https://www.ncbi.nlm.nih.gov/geo/query/acc.cgi?acc=GSE129699). One gene, *CTR1*, was identified from the pooled library that did not express CYP1A2; this gene functions in high-affinity copper and iron transport (Dancis et al., 1994), and its human homolog confers drug resistance (Furukawa et al., 2008), indicating that *CTR1* is involved in xenobiotic transport. We speculate that higher AFB_1_ concentrations are required to identify additional resistance genes in the yeast library lacking metabolic activation.

Using the same stringent assessment (q < 0.1), in three independent screens, we identified 95 genes that confer AFB_1_ resistance in cells expressing CYP1A2, of which 85 genes have been ascribed a function. Of these, 15 DNA repair genes (18%) were identified (Table1) and another 70 genes have diverse functions (Table 2). The gene that scored the most highly significant was *RAD54*. Five highly significant genes, participating in nucleotide excision repair, DNA damage tolerance, and ribosome assembly were twice found. An additional five genes were found that that were highly significant (q < 0.1) in one screen and significant in another screen (p < 0.05). Among these were those involved in proteolysis (*CUE1*), vacuolar acidification (*VOA1*), cell cycle progression (*FKH2*), DNA recombinational repair (*RAD55*) and DNA damage tolerance *(RAD18*). Selected genes were confirmed by growth curves (Figure 2); AUCs for additional genes are listed in supplemental Figure 1.

**Figure 2:**
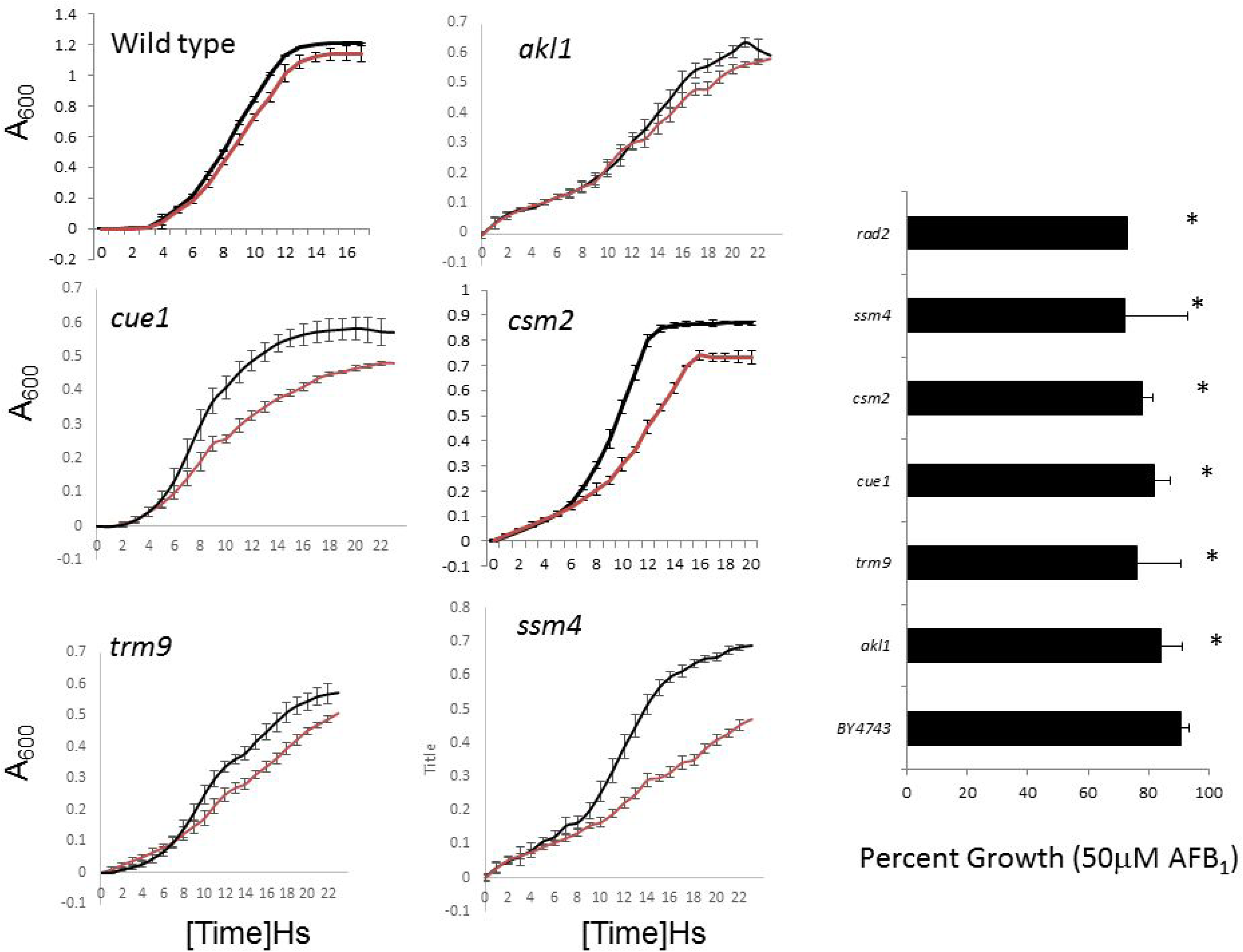
Growth curves for selected diploid mutants identified in the high throughput screen. A_600_ is plotted against time (Hs). Standard deviations are indicated at 1 h time points. The growth curves are indicated for wild type (BY4743) and *csm2*, *alk1*, *ssm4*, *cue1*, and *trm9*. The bar graph indicates area % growth of AFB_1_-exposed strains as determined by ratio of the area under curves (area under the curve for treated strain/area under the curve for strain exposed to DMSO x 100%).

**Table 1.**
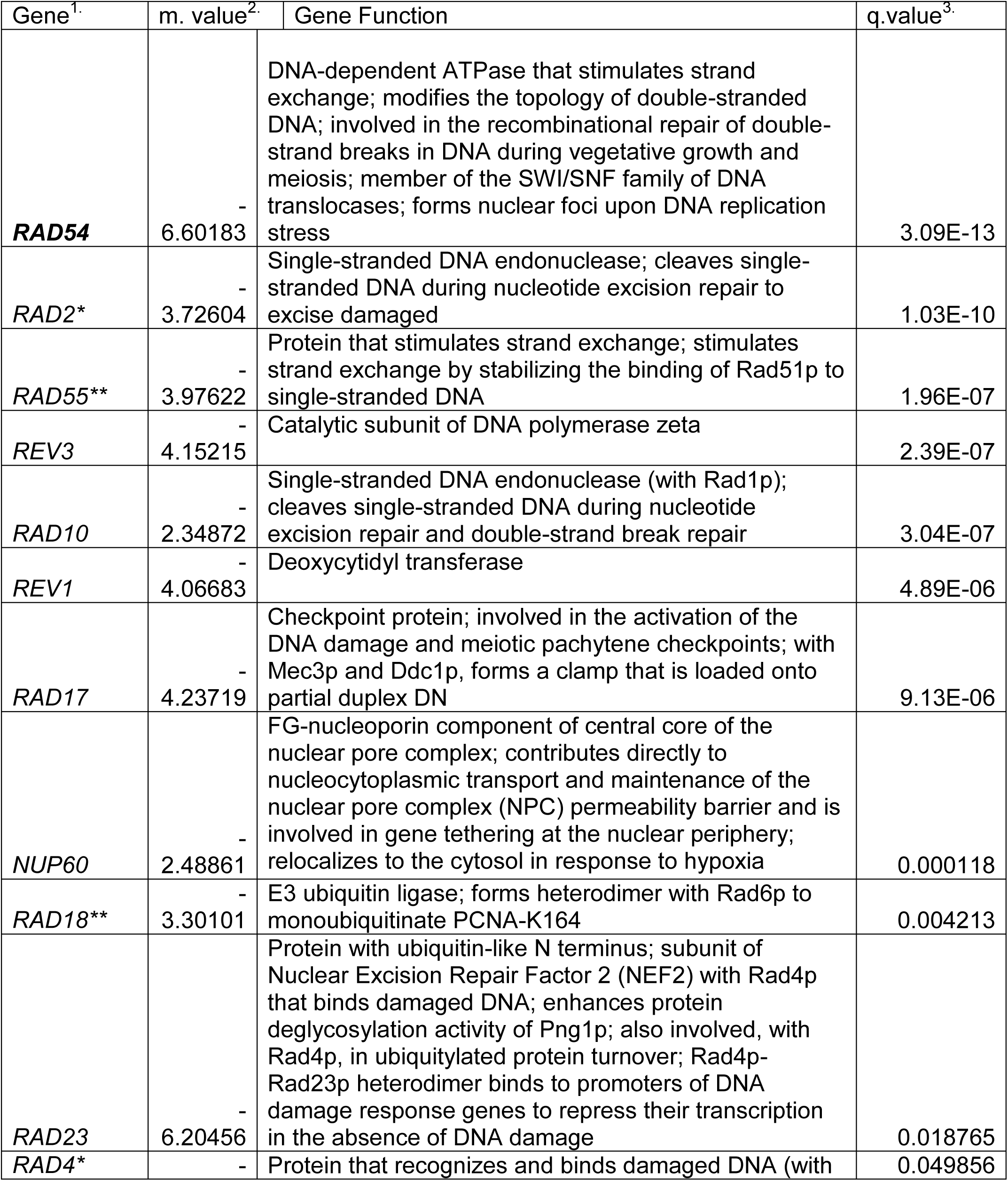
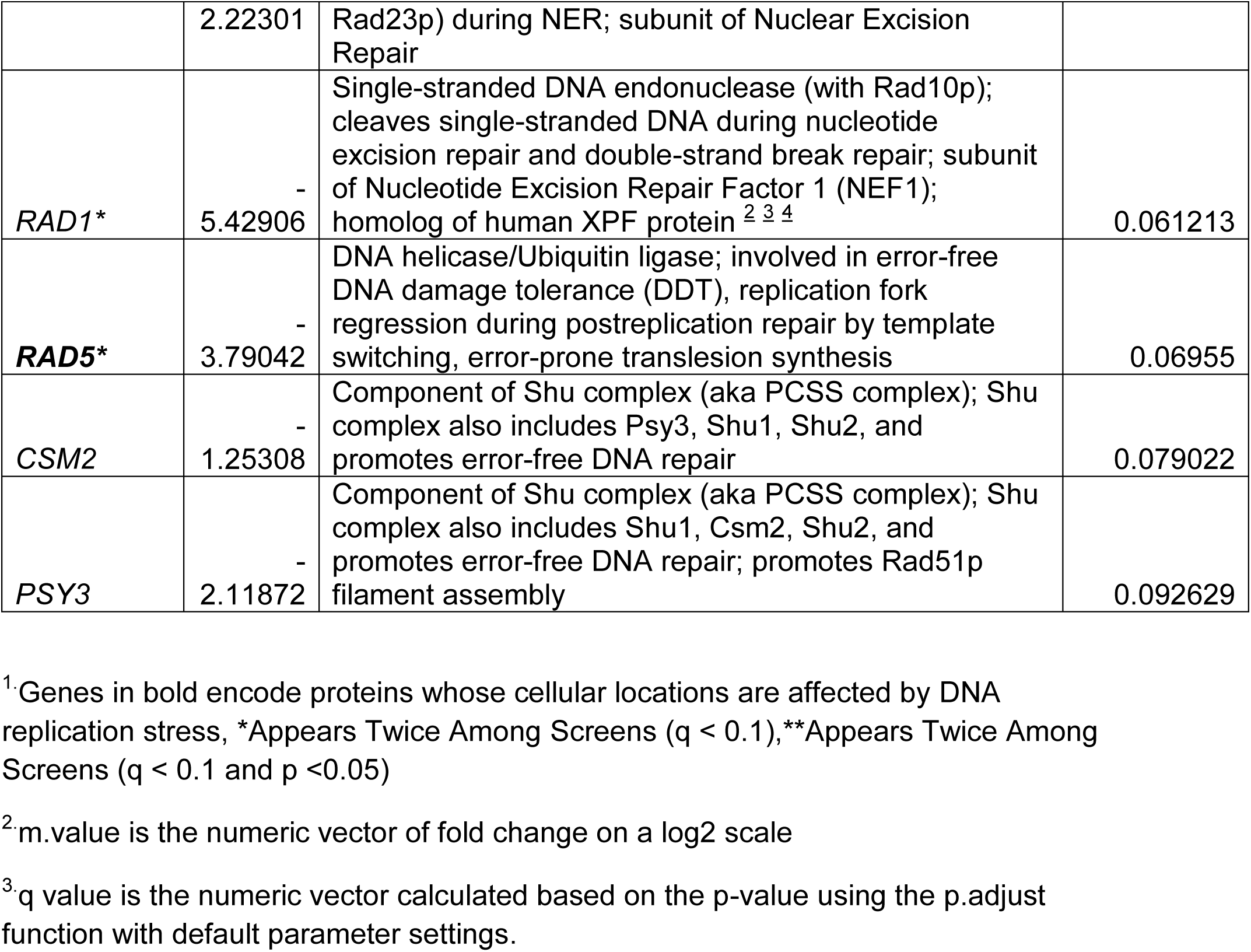
Fitness Scores for 15 AFB_1_ Resistant Genes Related to DNA Repair and Ranked by Significance

**Table 2.**
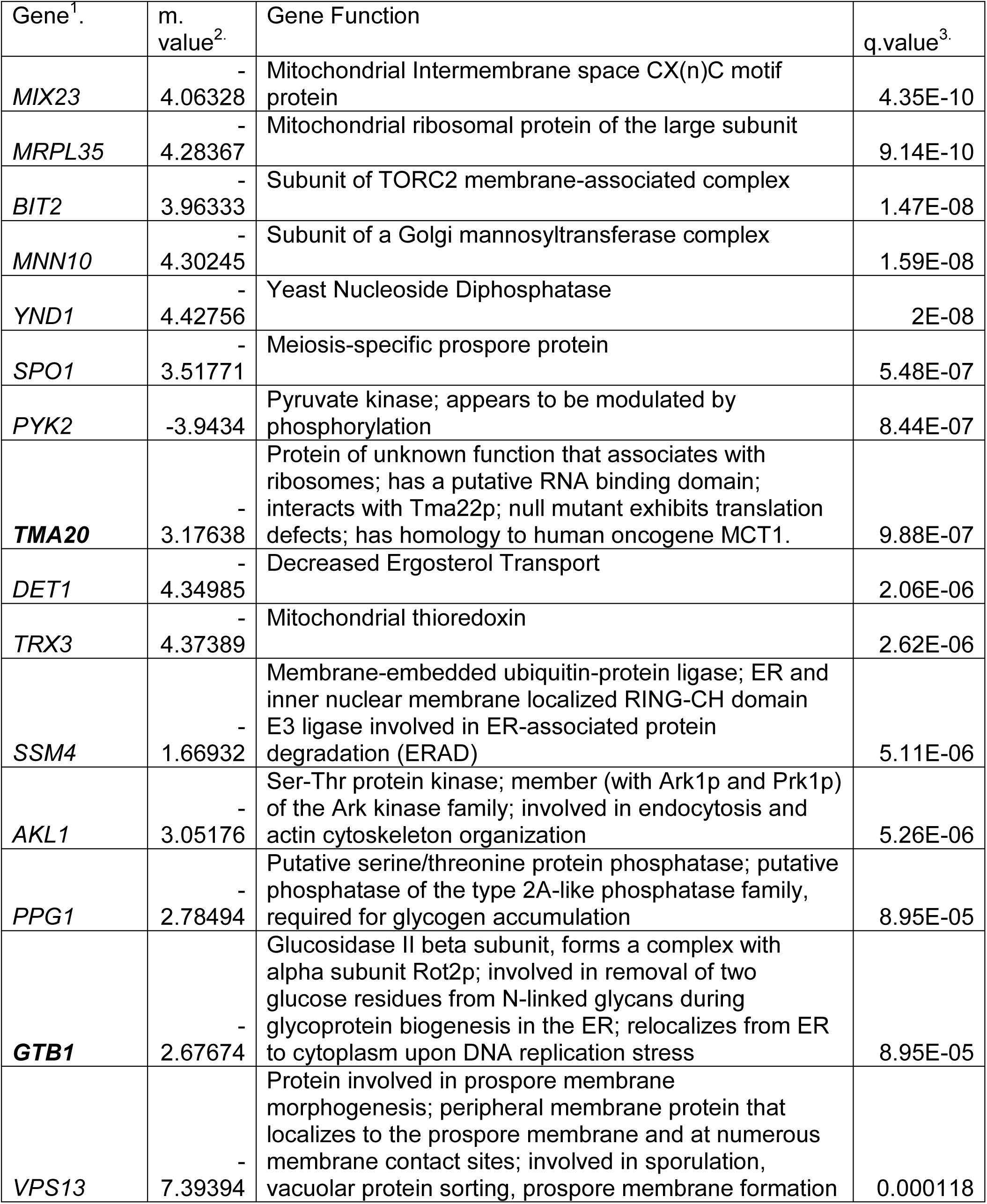
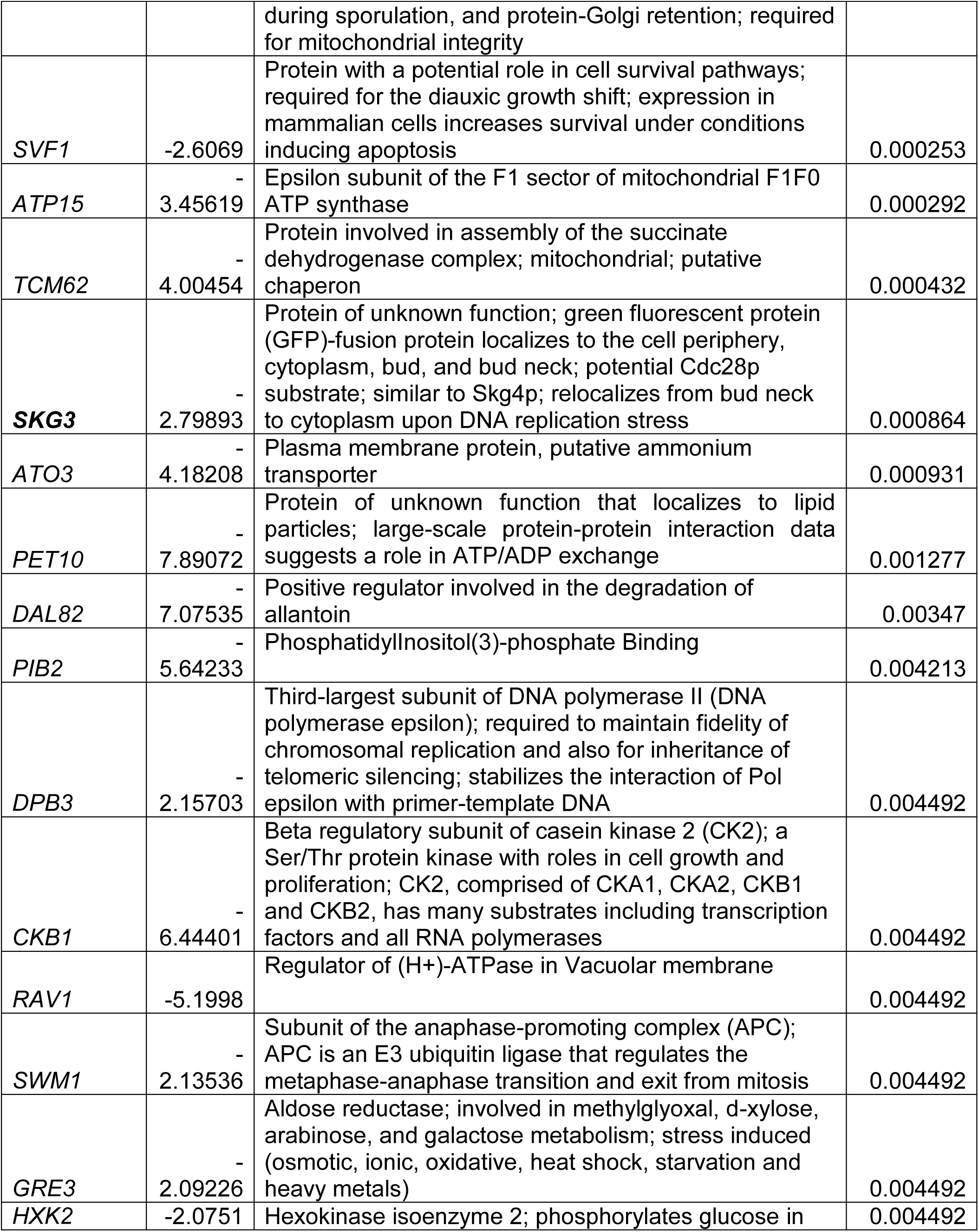
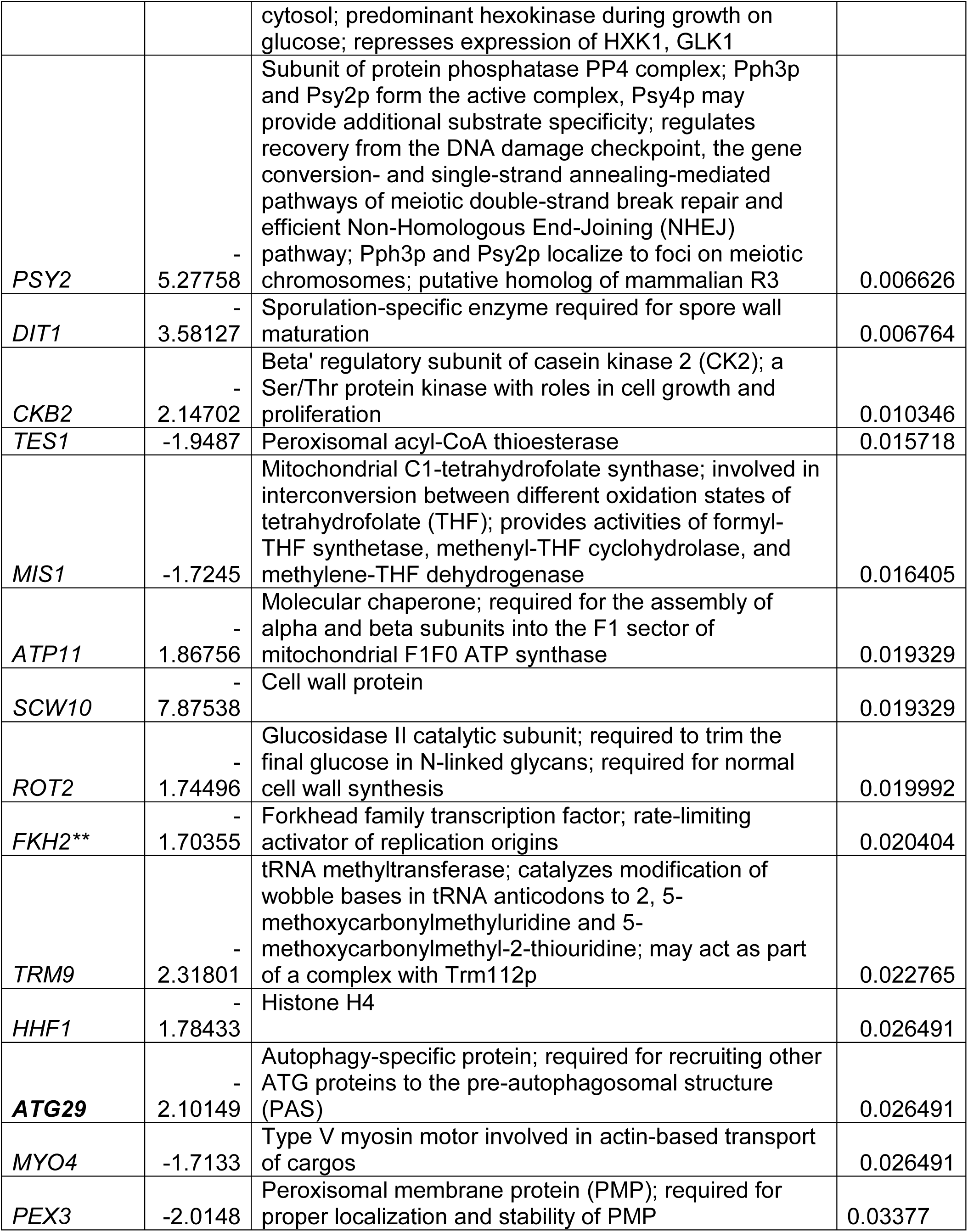
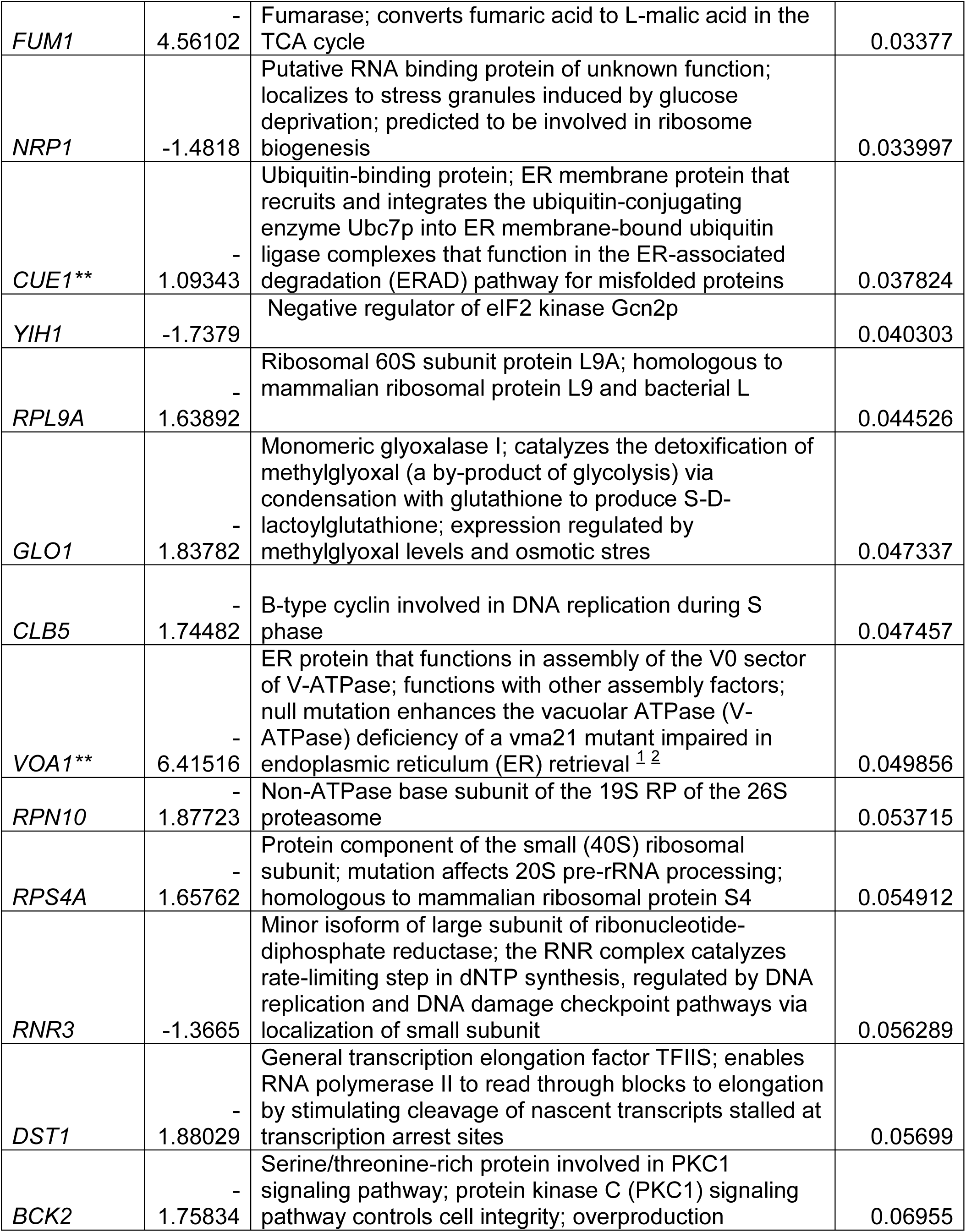
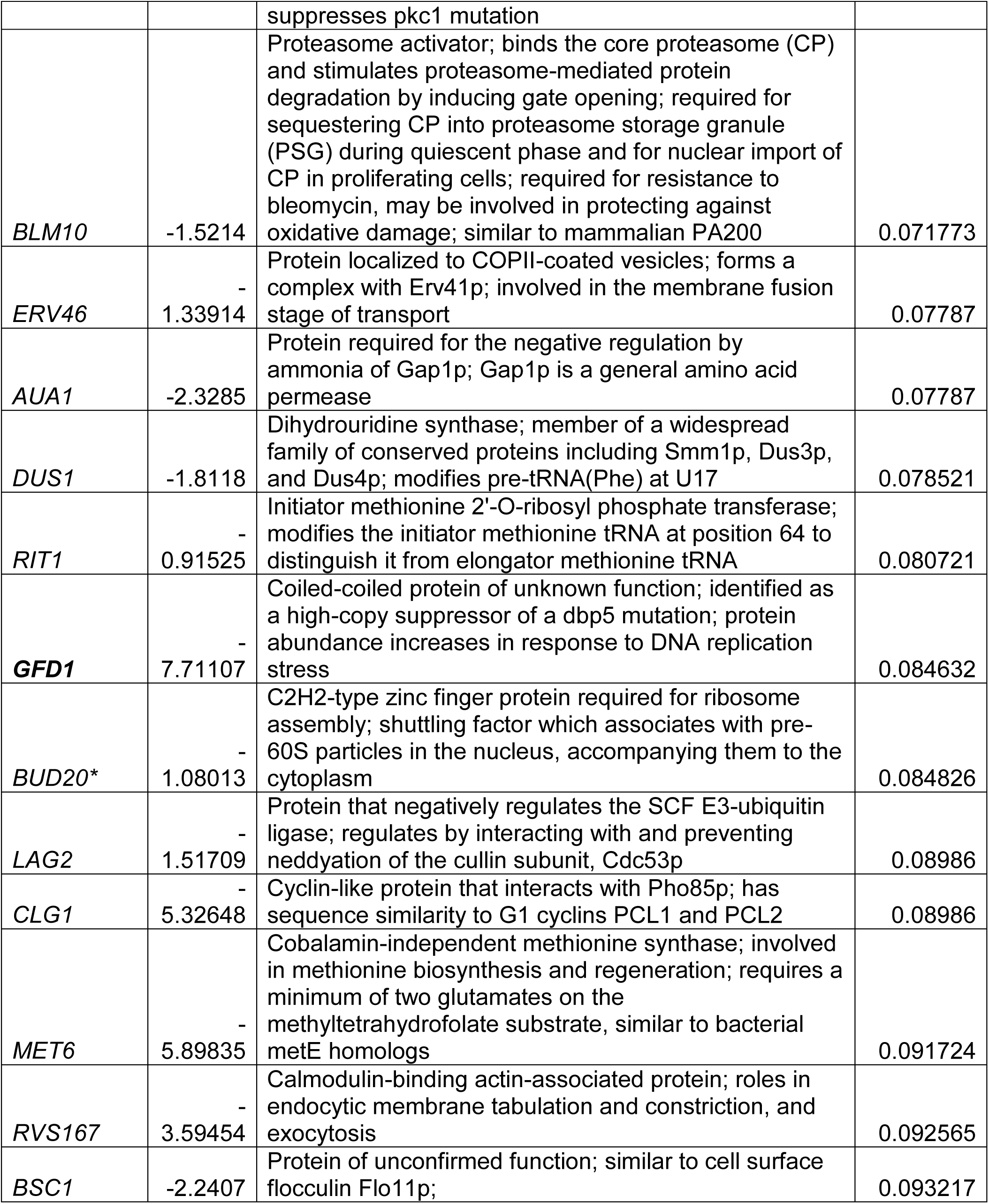
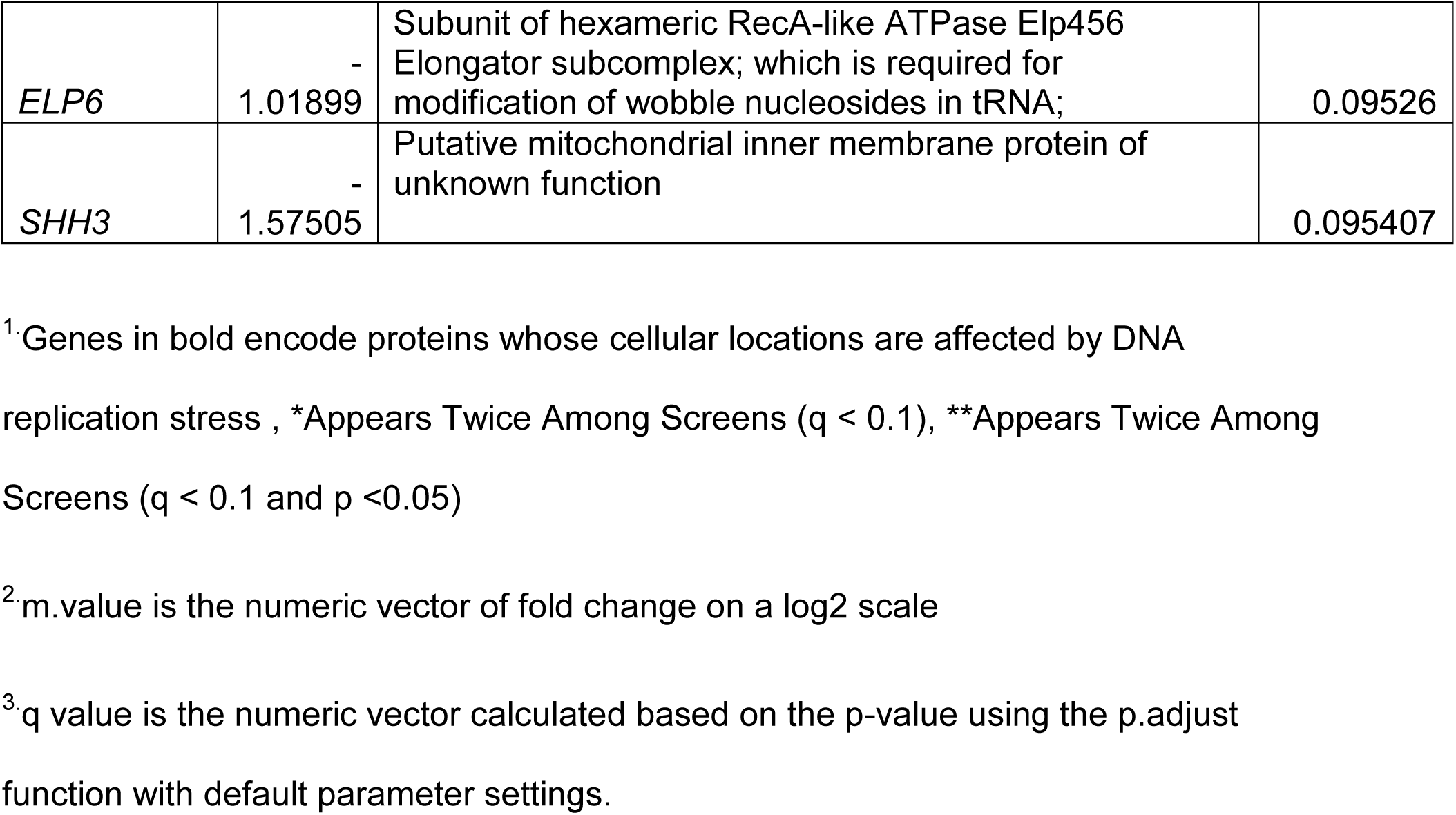
Fitness Scores for 70 AFB_1_ resistant genes ranked by significance

According Yeast GO-Slim Process and Funspec MIPS GO grouping, resistance genes included those that function in the DNA damage response, DNA repair, recombination, and DNA damage stimulus, tRNA modifications, carbohydrate metabolism, and cell cycle progression; the top fifteen GO groups are shown in Table 3 (for full list, see supplemental Table 3). Carbohydrate metabolism genes that function in cell wall maintenance and glycogen metabolism were previously identified to confer resistance to a variety of toxins, such as benzopyrene and mycophenolic acid (O’Connor et al., 2012). Other genes involved in carbohydrate metabolism, such as *GRE3* that encodes aldose reductase, could have a direct role in detoxification and is induced by cell stress (Barski et al., 2008). Genes involved in rearrangement of the cellular architecture include *BIT2*, *AKL1* and *PPG1*; these genes function to rearrange the cellular architecture when cells are stressed (Schmidt et al., 1996). Thus, among AFB_1_ resistant genes are those that function to maintain structural integrity by affecting the cytoskeletal and cell wall architecture.

**Table 3.**
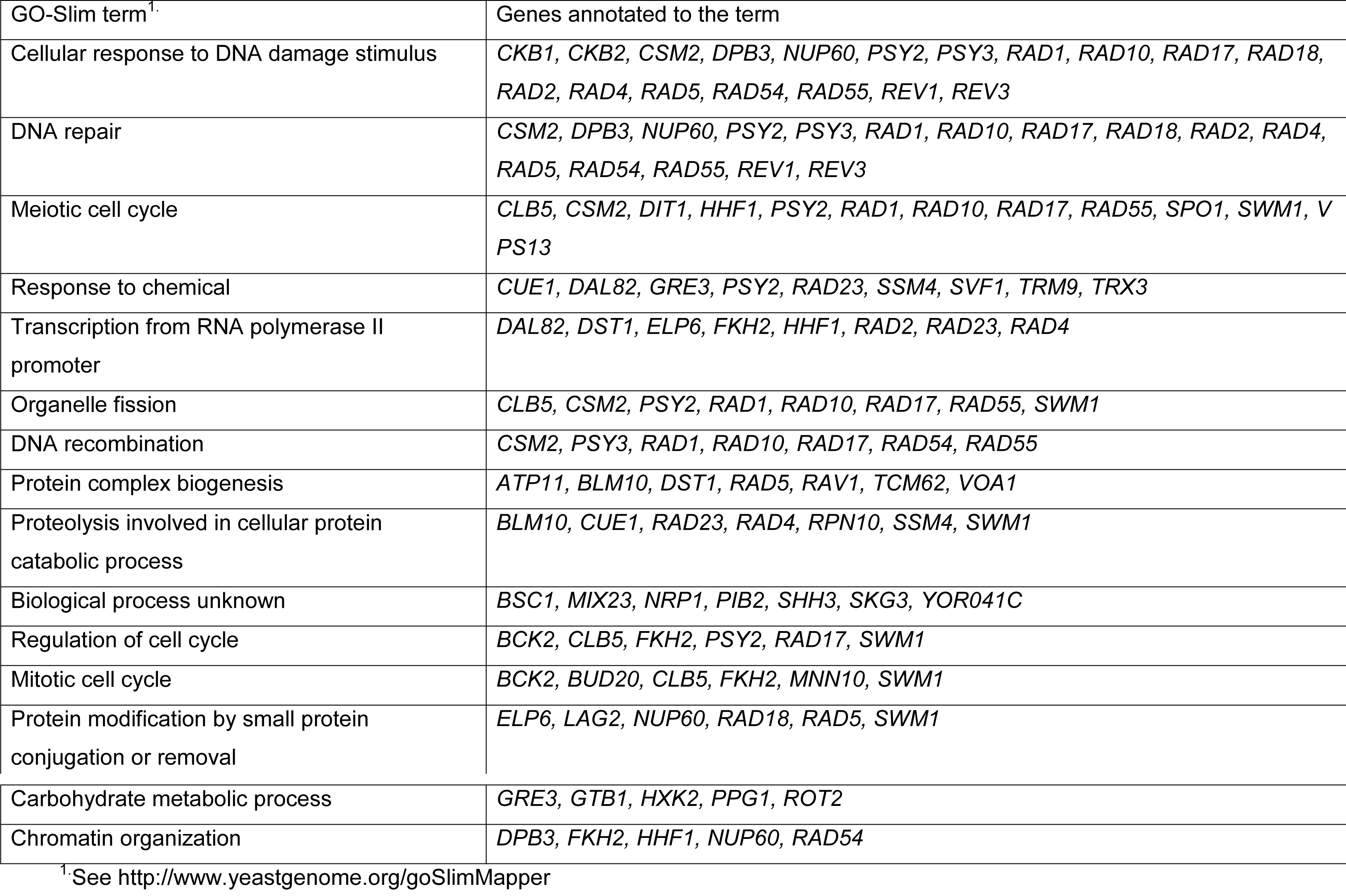
Top 15 Gene Ontology Groups According to the Yeast GO-Slim Process

| GO-Slim term<sup>1</sup>. | Genes annotated to the term |
| --- | --- |
| Cellular response to DNA damage stimulus | *CKB1, CKB2, CSM2, DPB3, NUP60, PSY2, PSY3, RAD1, RAD10, RAD17, RAD18, RAD2, RAD4, RAD5, RAD54, RAD55, REV1, REV3* |
| DNA repair | *CSM2, DPB3, NUP60, PSY2, PSY3, RAD1, RAD10, RAD17, RAD18, RAD2, RAD4, RAD5, RAD54, RAD55, REV1, REV3* |
| Meiotic cell cycle | *CLB5, CSM2, DIT1, HHF1, PSY2, RAD1, RAD10, RAD17, RAD55, SPO1, SWM1, VPS13* |
| Response to chemical | *CUE1, DAL82, GRE3, PSY2, RAD23, SSM4, SVF1, TRM9, TRX3* |
| Transcription from RNA polymerase II promoter | *DAL82, DST1, ELP6, FKH2, HHF1, RAD2, RAD23, RAD4* |
| Organelle fission | *CLB5, CSM2, PSY2, RAD1, RAD10, RAD17, RAD55, SWM1* |
| DNA recombination | *CSM2, PSY3, RAD1, RAD10, RAD17, RAD54, RAD55* |
| Protein complex biogenesis | *ATP11, BLM10, DST1, RAD5, RAV1, TCM62, VOA1* |
| Proteolysis involved in cellular protein catabolic process | *BLM10, CUE1, RAD23, RAD4, RPN10, SSM4, SWM1* |
| Biological process unknown | *BSC1, MIX23, NRP1, PIB2, SHH3, SKG3, YOR041C* |
| Regulation of cell cycle | *BCK2, CLB5, FKH2, PSY2, RAD17, SWM1* |
| Mitotic cell cycle | *BCK2, BUD20, CLB5, FKH2, MNN10, SWM1* |
| Protein modification by small protein conjugation or removal | *ELP6, LAG2, NUP60, RAD18, RAD5, SWM1* |
| Carbohydrate metabolic process | *GRE3, GTB1, HXK2, PPG1, ROT2* |
| Chromatin organization | *DPB3, FKH2, HHF1, NUP60, RAD54* |

<sup>1</sup>See http://www.yeastgenome.org/goSlimMapper

Other gene ontology groups encompass mitochondrial maintenance and response to oxidative stress, and RNA metabolism and translation (supplemental Table 3). Genes involved in mitochondrial function and response to oxidative stress include *TRX3*, *MRPL35*, *MIX23*, *MIS1*; *TRX3* (thioredoxin reductase) functions to reduce oxidative stress in the mitochondria (Greetham and Grant, 2009). RNA metabolism genes include those that function tRNA modifications, including *MIS1, TRM9, DUS1,* and *RIT1* and RNA translation, such as *TMA20*. *TRM9* confers resistance to alkylated DNA damage, and links translation with the DNA damage response (Begley et al., 2007). These genes are consistent with the notion that AFB_1_ causes oxidative damage and that mitochondria are targets of AFB_1_-induced DNA damage.

In grouping genes according to cellular components (Yeast Mine), DNA repair complex was readily identified. Among the DNA repair complex was the Shu complex, and the NER complex I and II complex (Table 4). However, other interesting complexes that were identified included the glycosidase II complex, and the CK2 complex. Because DNA repair and DNA damage response genes were most prominent of the GO groups, we focused on the function of these genes in conferring AFB_1_ resistance. As expected, these genes included those in DNA recombination (*RAD54, RAD55*), nucleotide excision repair (*RAD1, RAD4, RAD10*), and DNA damage tolerance (*RAD5, RAD18, REV1, REV3*). Many of these genes function in DNA damage tolerance both by directly interacting in the pathway and by affecting cell cycle progression. For example, *PSY2* and *CKB2* (Toczyski et al., 1997) promote cell cycle progression after cell cycle delay or arrest caused by stalled forks or double-strand breaks, respectively.

**Table 4.**
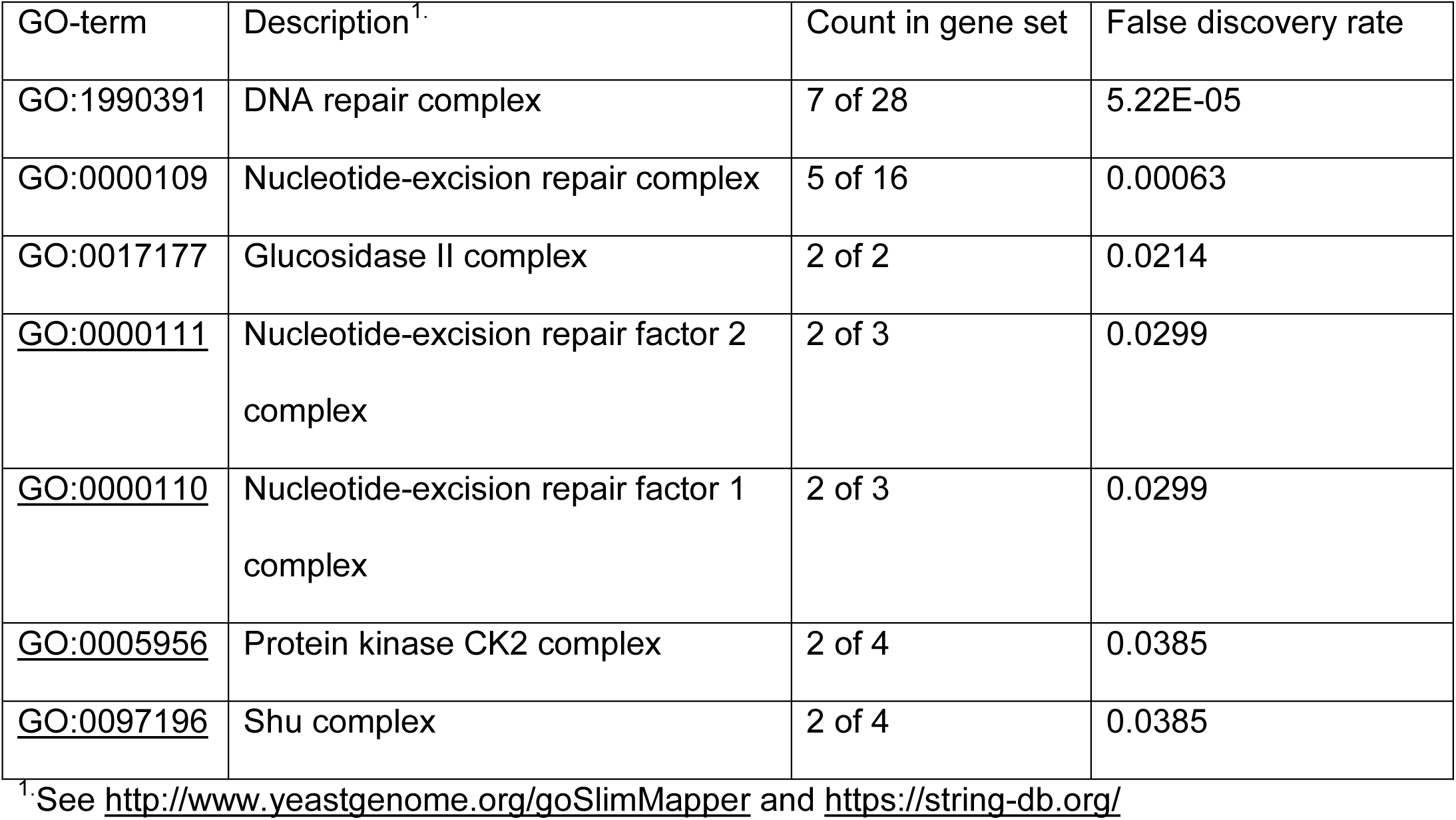
Protein Complexes That Participate in AFB_1_ Resistance

| GO-term | Description<sup>1</sup> | Count in gene set | False discovery rate |
| --- | --- | --- | --- |
| GO:1990391 | DNA repair complex | 7 of 28 | 5.22E-05 |
| GO:0000109 | Nucleotide-excision repair complex | 5 of 16 | 0.00063 |
| GO:0017177 | Glucosidase II complex | 2 of 2 | 0.0214 |
| <u>GO:0000111</u> | Nucleotide-excision repair factor 2 complex | 2 of 3 | 0.0299 |
| <u>GO:0000110</u> | Nucleotide-excision repair factor 1 complex | 2 of 3 | 0.0299 |
| <u>GO:0005956</u> | Protein kinase CK2 complex | 2 of 4 | 0.0385 |
| <u>GO:0097196</u> | Shu complex | 2 of 4 | 0.0385 |

<sup>1</sup>See http://www.yeastgenome.org/goSlimMapper and https://string-db.org/

To determine which GO biological processes and protein functions were most enriched among the AFB_1_ resistant genes we used the Panther software (Mi et al., 2016). Approximately 75% of the AFB_1_ resistant genes were involved in response to stress (Figure 3). A significant fraction of these genes also participate in DNA damage tolerance, including DNA post-replication repair and error-free and error-prone replication (Table 5). In classifying protein functions, we analyzed whether hydrolases, nucleases, phosphatases, DNA damage binding were more enriched among the 85 AFB_1_ resistant genes, compared to the genome, at large (Figure 3). Of these groups, only DNA damage binding demonstrated a significant difference (P < 0.05).

**Figure 3.**
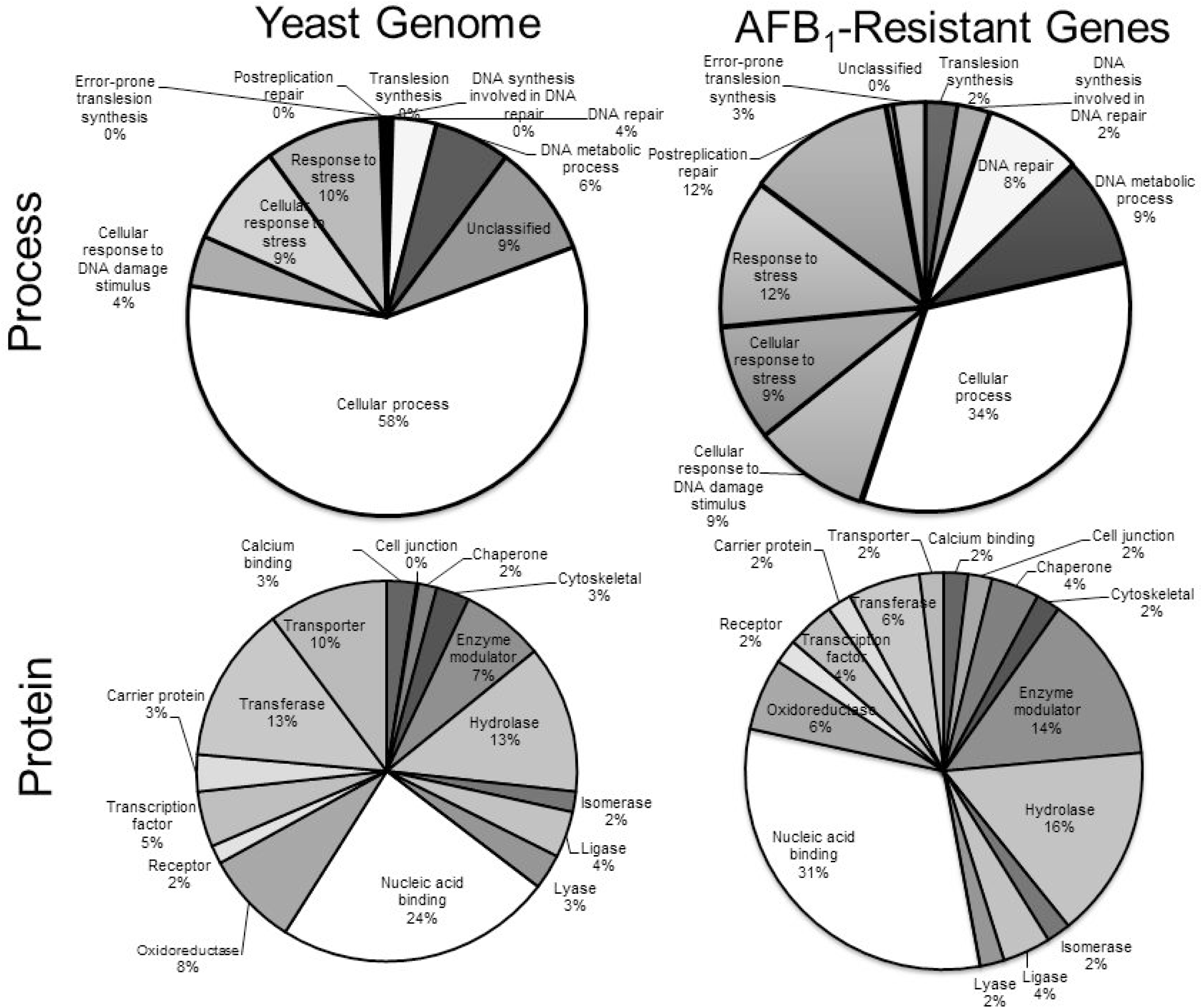
GO enrichment for the yeast genome compared to AFB_1_ resistant genes, as performed by Panther analysis. The top circles represent the % genes of that are grouped according to Process. The bottom group are those which are grouped according to protein class. GO groups included post replication repair, response to chemical, DNA repair, error-prone DNA synthesis, non-error prone DNA synthesis, and non-classified. Number of genes belonging to each GO is indicated within the pie.

**Table 5.** Enrichment of DNA Damage Repair Genes Among AFB_1_ Resistant Genes

| GO Biological Process<sup>1.</sup> | Yeast | AFB<sub>1</sub> Resistant Genes | Expected | Fold Enrichment | P value | Significance |
| --- | --- | --- | --- | --- | --- | --- |
| Error-prone translesion synthesis | 11 | 5 | <1 | >5 | 3.62E-04 | + |
| Translesion synthesis | 18 | 5 | <1 | >5 | 4.00E-03 | + |
| DNA synthesis involved in DNA repair | 20 | 5 | <1 | >5 | 6.67E-03 | + |
| DNA repair | 285 | 15 | 3.10 | 4.85 | 8.12E-04 | + |
| DNA metabolic process | 495 | 17 | 5.38 | 3.16 | 3.99E-02 | + |
| Cellular process | 4675 | 66 | 50.78 | 1.30 | 4.53E-02 | + |
| Cellular response to DNA damage stimulus | 331 | 18 | 3.60 | 5.01 | 2.63E-05 | + |
| Cellular response to stress | 673 | 23 | 7.31 | 3.15 | 8.74E-04 | + |
| Response to stress | 764 | 23 | 8.30 | 2.77 | 7.92E-03 | + |
| Postreplication repair | 29 | 5 | <1 | >5 | 3.96E-02 | + |
| Error-prone translesion synthesis | 11 | 5 | <1 | >5 | 3.62E-04 | + |
| Translesion synthesis | 18 | 5 | <1 | >5 | 4.00E-03 | + |
| DNA synthesis involved in DNA repair | 20 | 5 | <1 | >5 | 6.67E-03 | + |
| Unclassified | 733 | 1 | 7.96 | < 0.2 | 0.00E00 | - |

<sup>1.</sup> See http://pantherdb.org/tools/ for statistical analysis

To further determine the strength of the interactions among the AFB_1_ resistance genes, we performed interactome mapping, using STRING software (https://string-db.org/, Szklarczyk et al., 2019), which associates proteins according to binding, catalysis, literature-based, and unspecified interactions (Figure 4). The interactome complex in yeast included 86 nodes and 162 edges with a 3.77 average node degree. Besides the NER complexes, Individual complexes included the Shu complex, the Glucosidase II complex, and the protein kinase CK2 complex (Table 4); the glucosidase II complex is conserved in mammalian cells (Figure 4), While the strength and number of these interactions was particularly strong among the DNA repair genes, other interactions were elucidated, such as the interactions of protease proteins with cell cycle transcription factors and cyclins (Figure 4).

**Figure 4.**
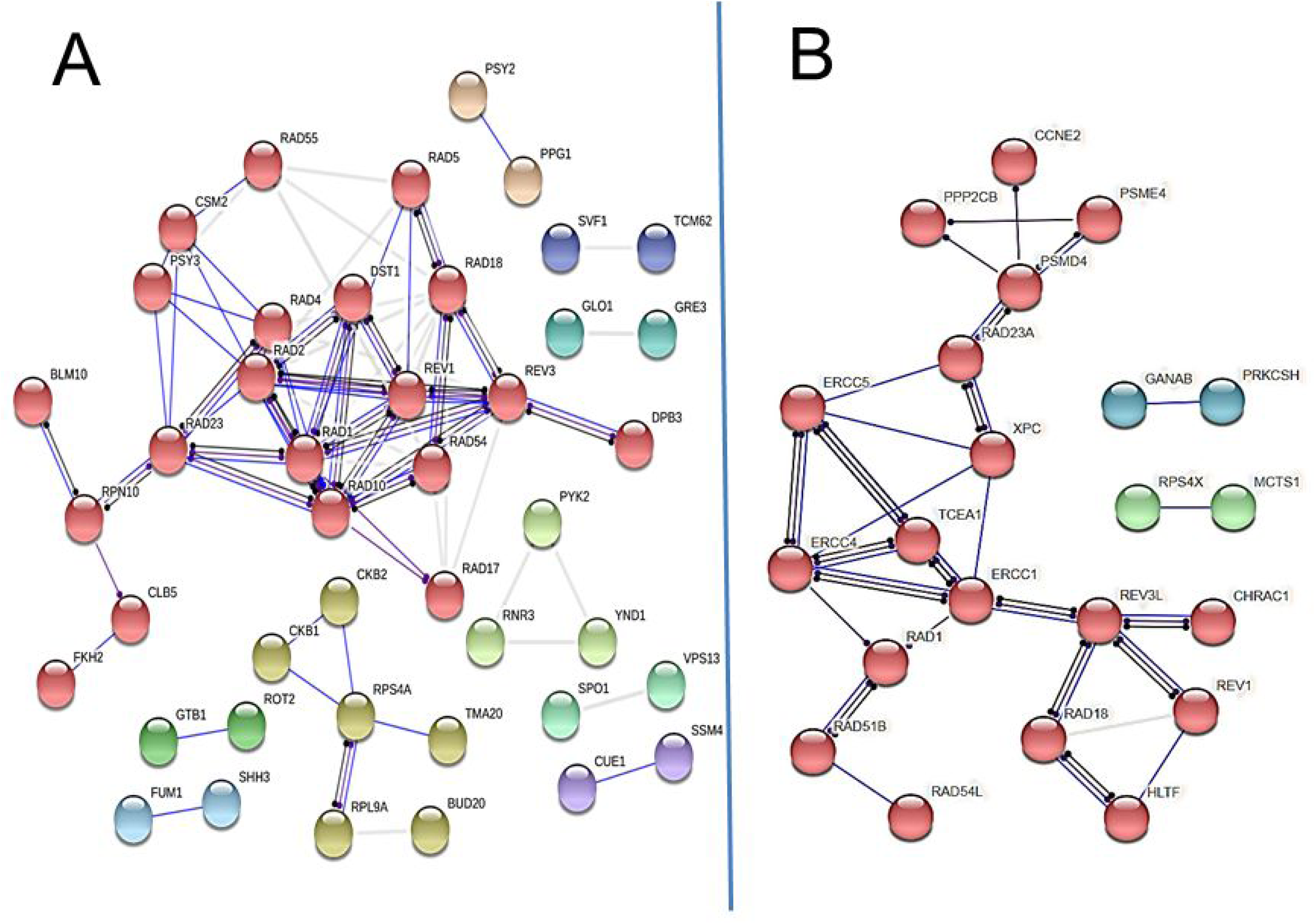
The protein interactome encoded by AFB_1_ resistance genes in budding yeast (left, A) and protein interactome encoded by their associated human homologs (right, B). The interactome was curated using String V11 (https://string-db.org, Szklarczyk et al., 2019), using a high confidence level of .8 and MCL cluster factor of 1.1. Proteins are represented by colored circles (nodes); different colors represent distinct interacting clusters. A core group, in red seen in both images, includes proteins that function in DNA repair pathways, and interact with proteases, transcription and cell cycle factors.. Lines represent the edges; a solid blue line indicates a binding event, a dark line indicates a reaction, and a purple line indicates catalysis. The lighter lines indicate a strong connection, as deduced from the literature. Lines that terminate with a dot indicate an unspecified interaction, whether positive or negative.

Since many known AFB_1_ resistance genes, which function in DNA repair and DNA damage tolerance pathways, were not present among highly significant genes (q < 0.1), we also used a less stringent (p < 0.05) qualifier to identify potential AFB_1_ resistance genes. Among genes identified were additional members of the SHU complex, including *SHU1* and *SHU2*. These genes were confirmed by additional growth curves (supplemental Figure 2).

The SHU complex was previously identified as participating in error-free DNA damage tolerance and mutation avoidance (Shor et al., 2005; Xu et al, 2013). The complex confers resistance to alkylating agents, such as methyl methanesulfonate (MMS), and cross-linking agents, such as cisplatin, but not to UV and X-ray (Godin et al, 2015). We previously showed that while X-ray associated unequal SCE (SCE) was *RAD5*-independent (Fasullo and Sun, 2017), MMS and 4NQO-associated unequal SCE occurs by well-conserved *RAD5*-dependent mechanisms (Fasullo and Sun, 2017; Unk et al., 2010). We therefore postulated that the SHU complex suppresses AFB_1_-associated mutagenesis while promoting AFB_1_-associated template switching. We introduced pCS316 (CYP1A2) into both the haploid wild-type strain (YB204) and a *csm2* mutant (YB558, see supplemental Table 1) to measure frequencies of AFB_1_-associated unequal SCE and *can1* mutations. Our results showed that while we observed a three-fold increase in SCE after exposure to AFB_1_ in wild type strains, we observed less than a two-fold increase in sister chromatid exchange in the *csm2* mutant (Figure 5). However, we observed a net increase in AFB_1_-associated Can^R^ mutations in the in *csm2* mutant, compared to wild type (P < 0.05). Average survival was only slightly higher in the wild type (51%) than in the *csm2* mutant (49%), but not statistically different (*P*= 0.8, N =4). We suggest that similar to MMS-associated DNA lesions, *CSM2* functions to suppress AFB_1_-associated mutagenesis while promoting template switching of AFB_1_-associated DNA adducts.

**Figure 5.**
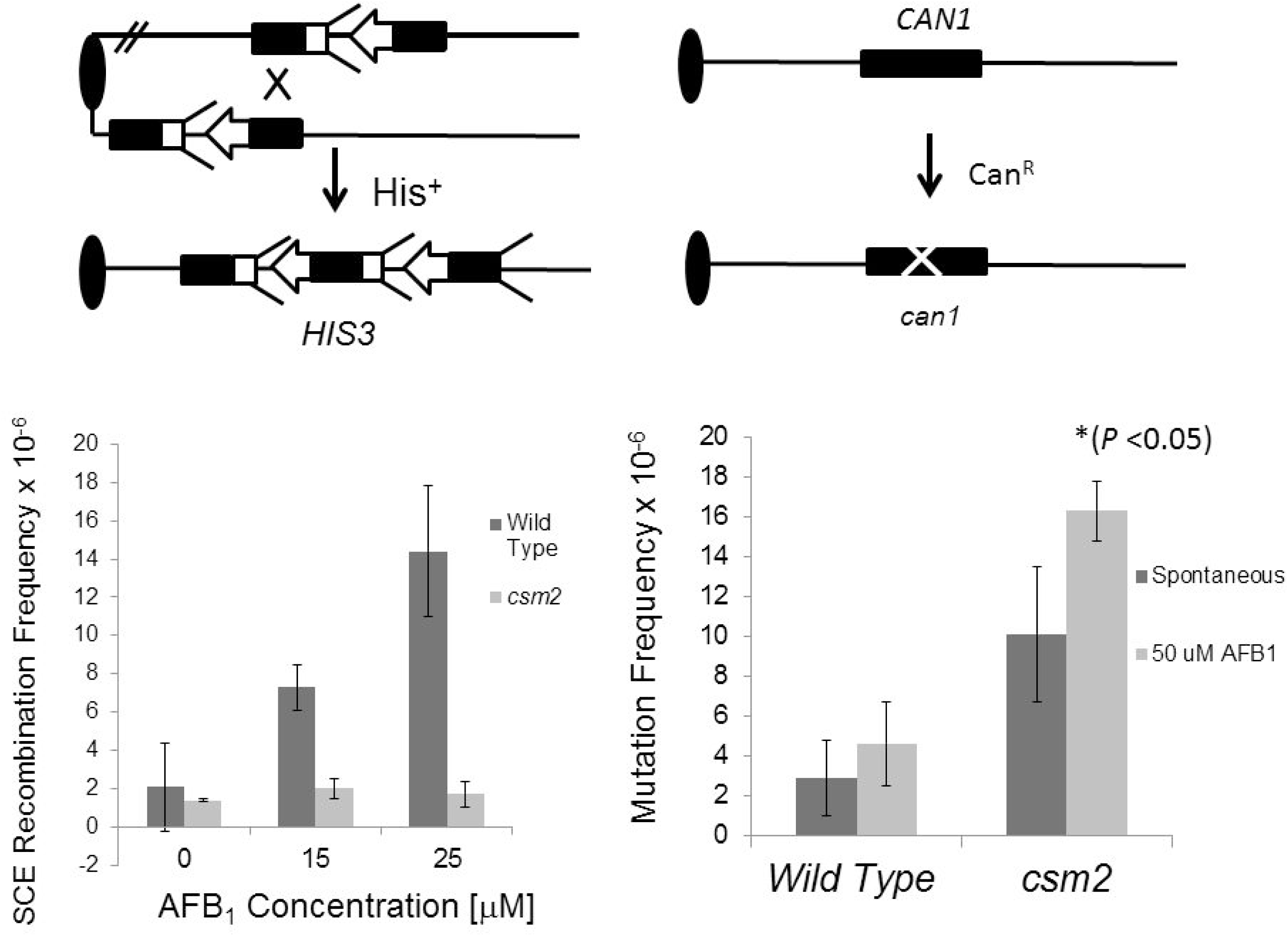
AFB_1_-associated sister chromatid recombination and mutagenesis frequencies in the wild type and *csm2* haploid mutant. The top part of the panel shows the assays for sister chromatid exchange and mutagenesis; for both assays the oval represents the centromere and the single line represents duplex DNA. For simplicity, the left arm of chromosomes IV and V are not shown. (Top left) Unequal sister chromatid recombination is monitored by selecting for His^+^ prototrophs that result from recombination between the juxtaposed, truncated *his3* fragments. The *his3*-Δ*3′* lacks the 3′ sequences (arrow head), while the *his3*-Δ*5′* lacks to promoter sequences (feathers). Both *his3* fragments are located with the amino acid reading frames oriented to the centromere. The *his3* fragments share a total of 450 bp sequence homology. (Top right). The mutation assay is measures the frequency of canavanine resistant mutants. The arrow notes the occurrence of point, missense, or deletion mutations that can occur in the *CAN1* gene.

If *CSM2* participates in a *RAD51*-dependent recombinational repair pathway to tolerate AFB1-associated DNA lesions, then we would expect that *RAD51* would be epistatic to *CSM2* for AFB1 resistance (Glassner and Mortimer, 1984). We measured AFB_1_ sensitivity in the *csm2*, *rad51* and *csm2 rad51* haploid mutants compared to wild type, using growth curves (Figure 6). Our data indicate the *csm2 rad51* double mutant is no more AFB_1_ sensitive compared to either *rad51* single mutants indicating that *CSM2* and *RAD51* are in the same epistasis group for AFB_1_ sensitivity. In contrast, *csm2 rad4* double mutants are more sensitive to AFB_1_ than either the *csm2* and *rad4* single mutants; the fitness measurement of the double mutant (0.071) is also less than the product of the *csm2* (0.34) and *rad4* (0.28) single mutants. These data indicate that *CSM2* participates in a *RAD51*-mediated pathway for AFB_1_ resistance, and similar to *RAD51*, confers AFB_1_ resistance in a *rad4* mutant (Fasullo et al., 2010).

**Figure 6.**
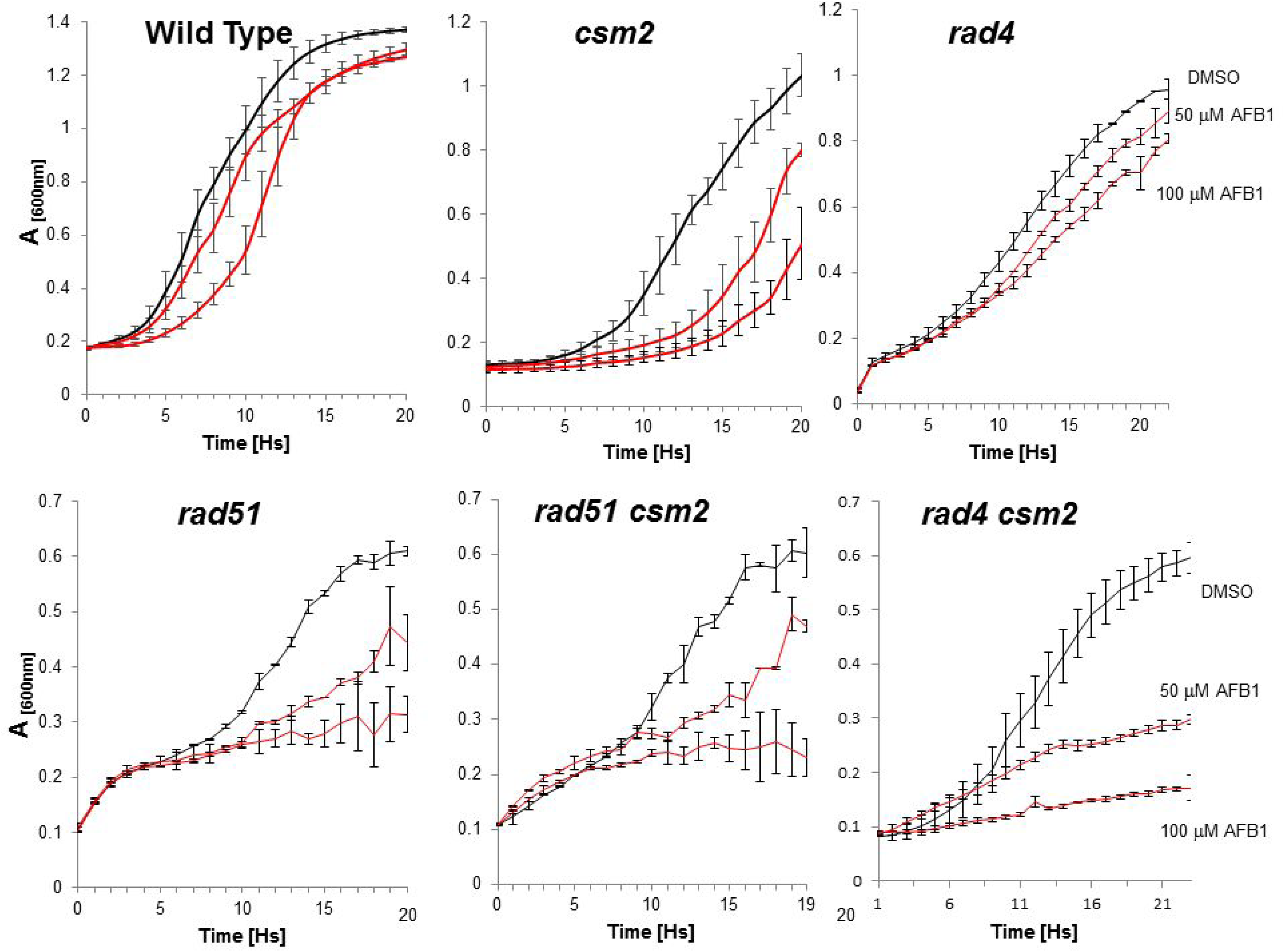
Growth of wild type (BY4741), *csm2*, *rad4*, *rad51*, *csm2 rad51*, and *csm2 rad4* cells after exposure to 50 and 100 μM AFB_1_. (Left) Growth of cells containing pCS316 and expressing CYP1A2 after chronic exposure to 0.5% and 1.0% DMSO (black), 50 *μ*M (red), and 100 *μ*M (red) AFB_1_. The relevant genotype is given above the panel (see Table 1, for complete genotype). Approximately 10^5^ log-phase cells were inoculated in each well, n *=* 2. *A*_600_ is plotted against time (hs). Bars indicate the standard deviations of measurements, n = 2.

Human orthologs for many essential yeast genes can directly complement the corresponding yeast genes (Kachroo et al., 2015). Human homologs are listed for 52 yeast AFB_1_-resistant genes. (Table 6). These homologs include those for DNA repair, DNA damage tolerance, cell cycle, and cell maintenance genes. Several of these genes, such as the human *CTR1* (Zhou and Gitschier, 1997), can directly complement the yeast gene. Other DNA human DNA repair genes, such as those that encode *RAD54* (Kanaar et al., 1996), *RAD5* (Unk et al, 2010), and *RAD10* orthologs, can partially complement sensitivity to DNA damaging agents (Aggarwal and Brosh, 2012).

**Table 6.**
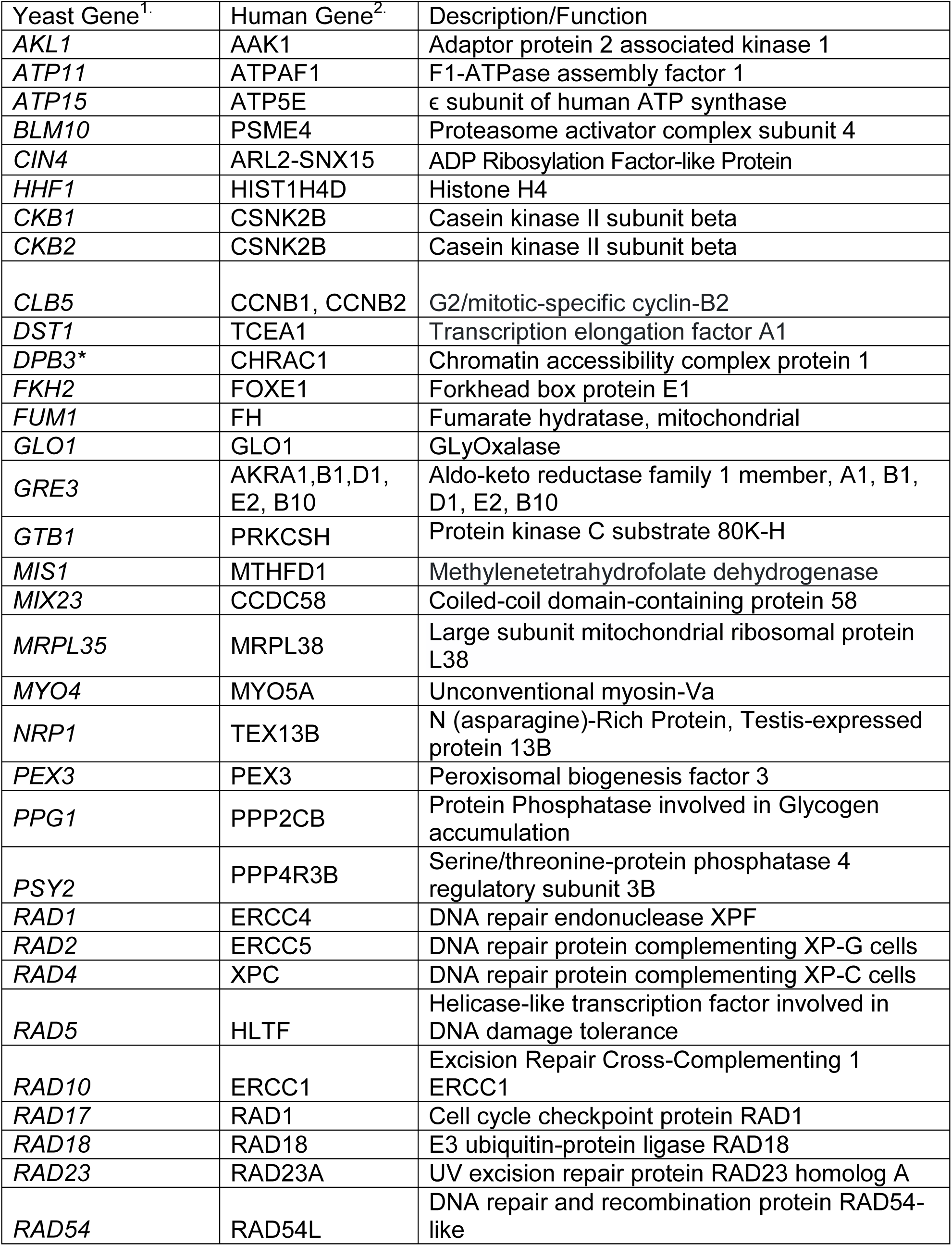
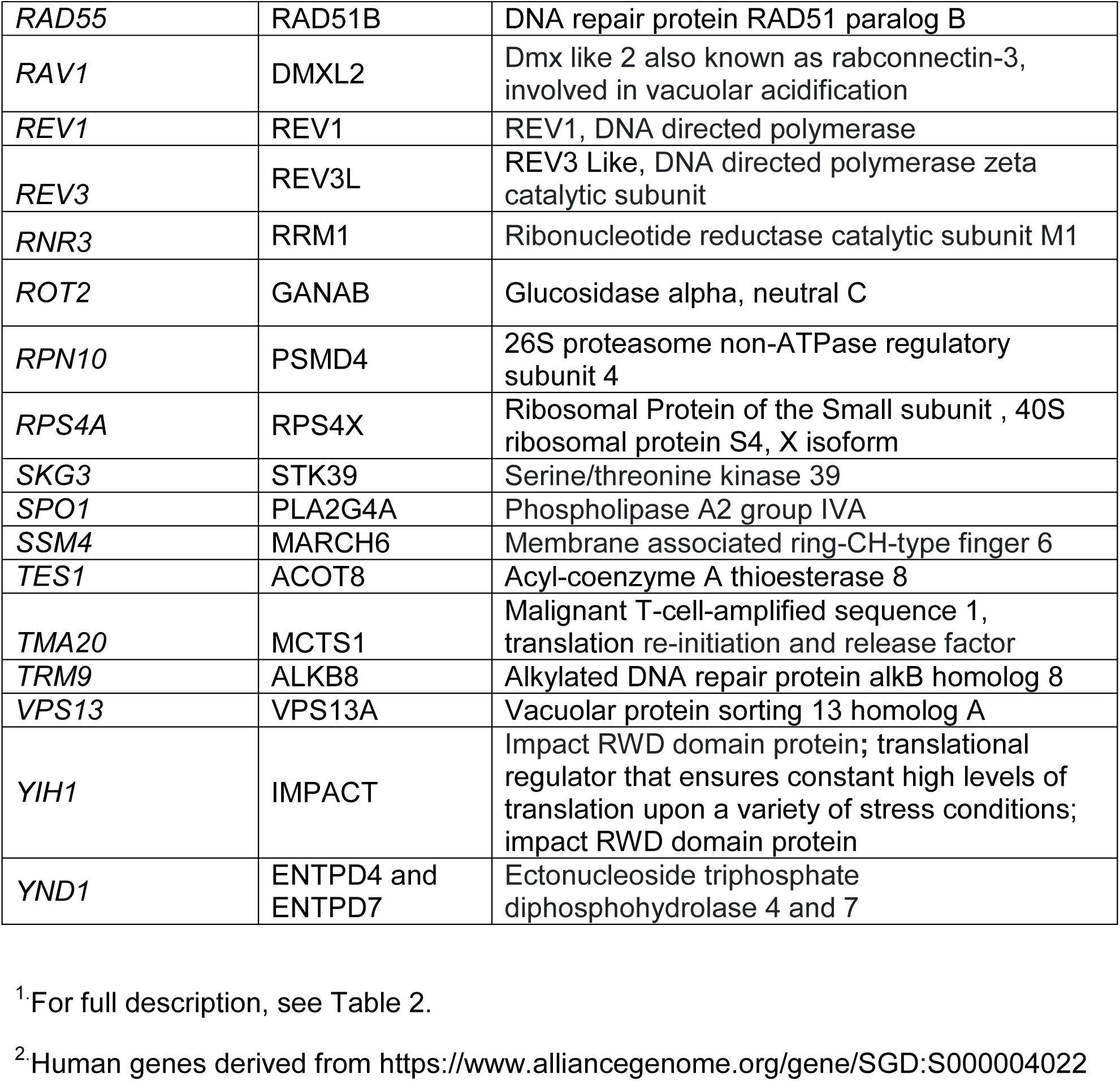
Human Genes Orthologous to Yeast Resistance Genes

## Discussion

Human CYP1A2-mediated activation of the mycotoxin AFB_1_ generates a highly reactive epoxide that interacts with DNA, RNA, and protein, forming adducts which interfere in replication, transcription, and protein function. Previous experiments have documented the role of checkpoint genes, RAD genes, and BER genes in conferring AFB_1_ resistance in budding yeast (Keller-Seitz et al., 2004; Guo et al., 2005; Fasullo et al, 2010). The goal of this project was to identify additional AFB_1_ resistance genes that may elucidate why AFB_1_ is a potent yeast recombinagen but a weak mutagen (Sengstag et al., 1996).

Here, we profiled the yeast genome for AFB_1_ resistance using three yeast non-essential diploid deletion libraries; one was the original library and the other two expressed human CYP1A2. We identified 96 resistance genes, of which 85 have been ascribed a function. These resistance genes reflect the broad range of functions, including cellular and metabolic processes, actin reorganization, mitochondrial responses, and DNA repair. Many of the DNA repair genes and checkpoint genes have been previously identified in screens for resistance to other toxins (Lee et al., 2005; Dela Rosa et al., 2017; Giaever and Nislow 2014). While, mitochondrial maintenance genes, and oxidative stress genes (Amici et al., 2007; Mary et al., 2012) are expected AFB_1_ resistant genes based on studies of individual mutants (Fasullo et al, 2010; Guo et al., 2005), we also identified novel resistance genes that participate in DNA damage tolerance, both by modulating checkpoint responses and by recombination-mediated mechanisms. Of key importance, the CSM2/SHU complex (Shor et al., 2005) was required for AFB_1_-associated sister chromatid recombination, underscoring the role of recombination-mediated template switch mechanisms for tolerating AFB_1_-associated DNA damage. Since many yeast genes are conserved in mammalian organisms (Bernstein et al, 2005), we suggest similar mechanisms for tolerating AFB_1_-associated DNA damage may be present in mammalian cells.

We used a novel reagent consisting of a pooled yeast library expressing human CYP1A2, on a multi-copied expression vector. Because CYP1A2 activates AFB_1_ when toxin concentration is low (Eaton and Gallagher, 1994), our modified yeast library mimicked AFB_1_ activation when the toxin is present in low concentrations in the liver. One limitation of the screen in the CYP1A2-expressing library is that AFB_1_-associated toxicity is not directly proportional to AFB_1_ concentration (Fasullo et al., 2010); we speculate that CYP1A2 activity is the limiting factor. Although individual yeast strains expressed similar amounts of CYP1A2 activity from among the subset of deletion strains tested, it is still possible that profiling resistance among individual deletion strains is influenced by the stability or variable expression of the membrane-associated human CYP1A2 (Murray and Sa-Correia, 2001). Secondly, many DNA repair and checkpoint genes that were previously documented to confer resistance, such as *RAD52*, were not identified in the screen (Fasullo et al, 2010, Guo et al, 2006). One possible reason is that some, such as *rad52,* grow poorly (Figure 1, Fasullo et al, 2008), and we suspect that other slow-growing strains dropped out early in the time course of exposure. Future experiments will more carefully assess generation times needed to detect known resistance genes in the library expressing CYP1A2.

Because metabolically activated AFB_1_ causes protein, RNA, and lipid damage (Weng et al., 2017), besides DNA damage, we expected to find a functionally diverse set of AFB_1_ resistance genes. Among AFB_1_ resistant genes were those involved in protein degradation and ammonia transport, actin reorganization, tRNA modifications, ribosome biogenesis and RNA translation. Some genes encoding these functions, such as *BIT2* and *TRM9,* have important roles in maintaining genetic stability and in double-strand break repair (Begley, et al., 2007; Schmidt et al., 1996; Schonbrun et al, 2012; Shimada et al., 2013).Several genes, such as *PPG1*, are involved in glycogen accumulation; these genes are also required to enter the quiescent state (Li et al., 2015). Other genes are involved in cell wall synthesis, including *MNN10*, *SCW10* and *ROT2*; we speculate that cell wall synthesis genes confer resistance by impeding AFB_1_ entrance into the cell, while genes involved in protein degradation in the ER may stabilize CYP1A2 and thus enhance AFB_1_-conferred genotoxicity (Murray and Sa-Correia, 2001). Glucan and other cell wall constituents have also been speculated to directly inactivate AFB1, and yeast fermentative products are supplemented in cattle feed to prophylactically reduce AFB1 toxicity (Pereyra et al., 2013).

Although AFB_1_-associated cellular damage is associated with oxidative stress (Amici et al., 2008; Liu and Wang, 2016; Mary et al., 2012) only a few yeast genes that confer resistance to reactive oxygen species (ROS) were identified in our screens. These genes included *TRX3*, *YND1*, *VPS13*, *BIT2, GTP1, FKH2, SHH3, NRP1,* and *BUD20*, which have a wide variety of functions (Bassler et al., 2012; Greetham and Grant, 2009; Schmidt et al., 1996). While known genes associated with oxidative stress and oxidative-associated DNA damage, such as *YAP1, SOD1*, and *APN1*, were not identified, other mitochondrial genes, such as *TRX3*, were identified. Guo et al. (2005) also showed that the haploid *apn1* mutant was not AFB_1_ sensitive. We offer two different explanations: first, the AFB_1_-associated oxidative damage is largely localized to the mitochondria, and second, there may be redundant pathways for conferring resistance to AFB_1_-associated oxidative damage, and therefore single genes were not identified. It is most likely the later as screens with oxidants (like t-BuOOH) also fail to identify expected antioxidant enzymes (Said et., al 2004).

A majority of the AFB_1_ resistant genes belong to GO groups that include response to chemical, response to replication stress, and post replication repair. Proteins encoding functionally diverse genes, such as *RAD54*, *RAD5, GFD1*, *TMA20*, *SKG3*, *GRE3*, and *ATG29* (Table 3), are repositioned in the yeast cells during DNA replication stress (Tkach et al., 2012). The requirement for unscheduled DNA synthesis was illustrated by identifying genes involved in the DNA damage-induced expression of ribonucleotide reductase; these included *RNR3* and *TRM9*. *TRM9*, involved in tRNA modification, functions to selectively translate DNA damage-inducible genes, such as *RNR1* (Begley et al, 2007).

One unifying theme was that cell cycle progression and recovery from checkpoint-mediated arrest is a prominent role in mediating toxin resistance. *FKH2* functions as a transcription factor that promotes cell cycle progression and G_2_-M progression. Other genes are involved in the modulation of the checkpoint response, such as *PSY2*. While *CKB1* and *CKB2* have broad functions, including histone phosphorylation and chromatin remodeling (Barz et al., 2003; Cheung et al., 2005), *CKB2* is also required for toleration of double-strand breaks (Toczyski et al., 1997; Guillemain et al., 2007), and thus may function for the toleration of AFB_1_-associated damage. These genes support the notion that some of the AFB_1_-associated DNA adducts are well tolerated and can be actively replicated.

While we expected to identify individual genes involved in postreplication repair, such as *RAD5, REV1, REV3,* the *CSM2/PSY3* complex is novel. Absence of *CSM2* confers deficient AFB_1_-associated SCE but higher frequencies of AFB_1_-associated mutations, suggesting that *CSM2* functions to suppress AFB_1_-associated mutations by *RAD51*-mediated template switch mechanisms (Fasullo et al., 2008; Guo et al., 2006). Consistent with this idea, *RAD51* is epistatic to *CSM2* in conferring AFB_1_ sensitivity, while *rad4 cms2* double mutant exhibits synergistic AFB_1_ sensitivity with respect to the single *rad4* and *csm2* single mutants. However, *rad4 rad51* mutants still exhibit more AFB_1_ sensitivity than *rad4 csm2*, suggesting that *RAD51* may be involved in conferring resistance to other AFB_1_-associated DNA lesions, such as double-strand breaks. We also expect that the *RAD51* paralogs, *RAD55* and *RAD57*, share similar AFB_1_-associated functions with *RAD51*. Considering that *RAD57* is the XRCC3 ortholog, determining whether yeast *RAD51* paralogs suppress AFB_1_-associated mutation will aid in identifying similar complexes in mammalian cells. Such complexes may elucidate why XRCC3 polymorphisms are risk factors in AFB_1_-associated liver cancer (Long et al., 2008; Ji et al., 2015)

Human homologs of several of the identified yeast genes (Table 6) are, hyper-methylated, mutated, or over-expressed in liver disease and cancer. For example, mutations and promoter methylations of the human *RAD5* ortholog, Helicase-Like Transcription Factor (HLTF), are observed in hepatocellular carcinoma (Zhang, 2013; Dhont, 2016). Mutations in PRKCSH and GANAB, human orthologs of *GTB1* and *ROT2*, are linked to polycystic liver disease (Porath et al., 2016; Perugorria and Banales, 2017). The *CKB2* ortholog, CSNK2B (Chua et al., 2017; Dotan et al., 2001; Zhang et al., 2015), is over-expressed in several liver cancers and therapeutics are currently in clinical trial (Gray et al., 2014, Li et al., 2017, Trembley et al., 2017). It is tempting to speculate that over-expression of CSNK2B also confers AFB_1_ resistance.

Human homologs of other yeast genes have been correlated to the progression of other cancers, including colon cancer. These include the human homolog for *TRM9*, ALKB8, and the human homolog for *TMA20*, MCT1, which can complement translation defects of *tma20* mutants and has been implicated in modulating stress (Herbert and Shi, 2001) and double-strand break repair (Hsu et al., 2007). MCT-1 overexpression and p53 is noted to lead to synergistic increases in chromosomal instability (Kasiappan et al., 2009).

In summary, we profiled the yeast genome for AFB_1_ resistance and identified novel genes that confer resistance. The novel genes included those involved in tRNA modifications, RNA translation, DNA repair, protein degradation, and actin reorganization. Genes that function in DNA damage response, post-replication repair, and DNA damage tolerance were over-represented, compared to the yeast genome. We suggest that the *CSM2* (SHU complex) functions to promote error-free replication of AFB_1_-asociated DNA damage, and it will be interesting to determine whether mammalian orthologs of the SHU complex function similarly.

## Acknowledgements

We acknowledge William Burhans for his encouragement and support, and grants from the National Institutes of Health: R21ES1954, F33ES021133, and R15ES023685-03. We would like to thank Chris Vulpe for his gift of the pooled BY4743 library and guidance, Mingseng Sun for initiating work on AFB_1_-induced recombination, Jonathan Bard for bioinformatics expertise, and Brian Kellner for technical contributions.

---

Supplemental Figure 1. Percent growth calculated by area under the curves (AUCs) for yeast strains containing designated deletions after exposure to 50 µM AFB and/or 100 µM AFB. Panel A: Percent growth calculated for specific deletion strains after exposure to 50 µM AFB. Panel B: Percent growth calculated for specific strains after exposure to 50 µM AFB and/or 100 µM AFB. Percent growth were calculated by determining the ratio of the AUC after exposure to toxin to the AUC exposed to solvent (dimethyl sulfoxide) alone.

Supplemental Figure 2. Growth curve of the diploid wild type (BY4743 + pCS316), *shu1* + pCS316, *shu2* + pCS316, and *psy3* + pCS316 after exposure to 50 µM AFB. Growth (A_600_) is plotted against time (Hs). Standard deviations are indicated at 1 h time points, n = 2. See supplemental figure 1 for percent growth.

